# Building plumbing influences the microdiversity and community assembly of the drinking water microbiome

**DOI:** 10.1101/2024.09.23.614467

**Authors:** Huanqi He, Linxuan Huo, Solize Oosthuizen-Vosloo, Kelsey J. Pieper, Aron Stubbins, Byungman Yoon, Ameet J. Pinto

## Abstract

Building plumbing microbial communities can significantly influence water quality at the point of use, particularly during periods of stagnation. Thus, a fine-scale understanding of factors governing community membership and structure, as well as environmental and ecological factors shaping building plumbing microbial communities is critical. In this study, we utilized full-length 16S ribosomal RNA (rRNA) gene sequencing to investigate the microdiversity and spatial-temporal dynamics of microbial communities in commercial and residential building plumbing systems. Bacterial operational taxonomic units (OTUs) within commercial buildings exhibited much lower microdiversity relative to the same OTUs in residential buildings. Microdiversity was associated with higher persistence and relative abundance of OTUs. Interestingly, amplicon sequencing variants within the same OTUs exhibited habitat preferences based on the building type while also demonstrating varying temporal turnover patterns. Dispersal limitation disproportionately governed community assembly in commercial buildings, whereas heterogeneous selection was the dominant ecological mechanism shaping the microbial community in residential buildings. Dispersal limitation in commercial buildings is consistent with larger building sizes and greater periods of water stagnation. Interestingly, the inability to explain the extent of heterogeneous selection-driven community assembly in residential locations using measured water chemistry may suggest a disproportionately large effect of fine-scale variation in plumbing characteristics on community assembly in residential locations.

## 1. Introduction

The configuration and design of building plumbing systems varies greatly depending on the building type (AWWA Premise Plumbing Committee, 2022; Schmidt et al., 2019). This can influence microbial community ecology and overall drinking water safety at both spatial (De Sotto et al., 2020; Persily et al., 2023) and temporal scales (AWWA Premise Plumbing Committee, 2022; Julien et al., 2020; Vosloo et al., 2023). These water quality impacts can be accelerated by stagnation (Ling et al., 2018; Montagnino et al., 2022; Proctor et al., 2020), plumbing architecture (Cruz et al., 2020; Ji et al., 2015; Lee et al., 2021; Schück et al., 2023), flow patterns (Julien et al., 2020; Liu and Reckhow, 2013), and biofilm formation (Ling et al., 2018; Yao et al., 2023). A key feature affecting water quality in building plumbing is stagnation. Unlike municipal drinking water distribution systems (DWDS), the water within building plumbing can experience extended periods of stagnation depending on the number of occupants and their water-use patterns (Logan-Jackson et al., 2023). Changes in residual disinfectant concentrations resulting from stagnation and temperature changes due to the mixing of hot and cold water are common factors impacting microbial communities in building plumbing (Ley et al., 2020; Proctor and Hammes, 2015). These impacts include microbial growth, reduction in microbial diversity (Greenwald et al., 2022), and accelerated biofilm formation (Collins et al., 2017; Hozalski et al., 2020; Ley et al., 2020). Though short-term stagnation is not uncommon (e.g., overnight, over weekends, or holidays), prolonged water stagnation (e.g., weeks to months) has been linked to the occurrence of opportunistic premise plumbing pathogens (OPPPs) and significant health risks (Aw et al., 2022; Dowdell et al., 2023; Hozalski et al., 2020; Huang et al., 2020; Logan-Jackson and Rose, 2021; Logan-Jackson et al., 2021; Moritz et al., 2010). Temporary flushing can be a preventative measure to reduce OPPP risks during low occupancy periods (Dowdell et al., 2023; Hozalski et al., 2020; Ley et al., 2020).

To date, most building plumbing investigations have utilized short-read sequencing techniques and OTU-based approaches to profile the drinking water microbiome. Short-read approaches can limit the discernable taxonomic resolution, while OTU-level clustering can mask fine-scale variation in microbial communities (Callahan et al., 2019, 2017). Different bacterial strains within the same species have repeatedly been shown to vary in their spatial and temporal distributions (Larkin and Martiny, 2017). Thus, closely related sub-taxa may partition across different niches within the same environment, a distinction that would typically be masked at coarser genetic resolution (Chase and Martiny, 2018). The microdiversity within species, also known as strain-level or sub-taxa diversity (Fodelianakis et al., 2022; Koeppel and Wu, 2013; Larkin and Martiny, 2017), has rarely been explored using full-length 16S rRNA gene sequencing for the drinking water microbiome. This leaves a significant knowledge gap regarding the finer-scale diversity and dynamics in the building plumbing ecosystem.

Perhaps more importantly, a focus on microdiversity could help uncover ecological processes that would otherwise be hidden at a coarser genetic resolution. Two major ecological processes are known to shape the spatio-temporal patterns in microbial communities: deterministic (niche-based) and stochastic (neutral) processes (Ayarza and Erijman, 2011; Chase and Myers, 2011; Stegen et al., 2013; Yuan et al., 2019). Deterministic mechanisms govern microbial populations via abiotic factors (environmental variables, e.g., temperature, pH, residual disinfectant) and biotic factors (interactions among species) owing to different habitat preferences and differential microbial fitness (Han et al., 2023). In contrast, the stochastic processes influence taxa loss and gain through random birth, death, and immigration events as well as through drift (Chase, 2010; Zhou et al., 2014). While OTU-based approaches indicate that stochastic processes primarily govern the building plumbing microbiome (Cai et al., 2023; Ke et al., 2024; Ling et al., 2018; Yang et al., 2024), other studies using ASV-based analysis have revealed a dominant role of deterministic forces (Kinnunen et al., 2017; Liu et al., 2024). These discrepancies could stem from (1) building-specific plumbing complexities that affect the microbial assembly, (2) the potential for certain ecological patterns to be overlooked by the OTU-level analysis, and (3) lower genetic resolution of short-read-based approaches.

Thus, the overall objective of this study is to investigate microdiversity within the context of building plumbing microbiomes across different building types. To accomplish this, we characterized microbial communities in tap water samples collected from both commercial buildings and residential households over a period of six months. We used full-length 16S rRNA gene sequencing and a combination of OTU- and ASV-based analyses to discern spatial-temporal turnover and habitat preferences of the building plumbing microbiome. In doing so, we also aim to uncover ecological processes that may drive the inter-population (i.e., strains) and intra-population (i.e., species) level diversity and community assembly.

## 2. Materials and methods

### 2.1 Sample collection and wet chemistry analysis

Drinking water samples were collected from three large commercial buildings (COM) and four inhabited residential households (RES) in Boston, MA, USA from June to November 2020. All buildings received chloraminated drinking water from the same DWDS. The sampled taps included RES kitchens (n = 3), COM kitchens (n = 3), RES bathrooms (n = 2), and COM laboratories (n = 3). Temperature, pH, conductivity, dissolved oxygen (DO), dissolved organic carbon (DOC), nitrogen species, and total chlorine were measured for each sample. All samples and the entire dataset used in this study were concurrently collected with our previous study (Vosloo et al., 2023), where detailed sampling processes and wet chemistry analyses are described.

### 2.2 DNA extractions, full-length 16S rRNA gene sequencing, and data processing

Drinking water samples (1500 mL/sample) were concentrated through enclosed Sterivex ™ filter units (0.22 um pore size, EMD Millipore, MA, USA). Genomic DNA was extracted from the filter membranes following a modified DNeasy PowerWater Kit® protocol (Vosloo et al., 2023). Full-length 16S rRNA gene sequencing was performed at the Georgia Genomics and Bioinformatics Core (University of Georgia, GA, USA) on the PacBio Sequel II platform (Pacific Biosciences, CA, USA). Details regarding the primers, polymerase chain reaction (PCR) conditions, library preparation, and generation of the circular consensus sequence (CCS) reads were described previously (Vosloo et al., 2023). Raw sequences were processed with DADA2 v. 1.30.0 (Callahan et al., 2019) in R v. 4.3.3 (R Core Team, 2013) to remove primers, denoise, and construct amplicon sequence variants (ASVs). The SILVA nr v.138.1 database was used for taxonomic assignment of ASVs and non-bacterial sequences were removed. Table S1 summarizes the reads per sample at different stages of data processing in DADA2.

### 2.3 Microdiversity, persistence, and variability

ASVs were clustered into OTUs with the VSEARCH algorithm (Rognes et al., 2016) at 98.7% sequence identity (Yarza et al., 2014). Here, OTUs are analogous to species while constituent ASVs (all ASVs clustered into an OTU) represent different strains or sub-taxa within a species. Microdiversity captures effective ASVs that contribute to the diversity within each OTU. For microdiversity calculation, we only considered OTUs with at least 5000 reads to ensure that an accurate Shannon diversity index can be captured with sufficient sequences. The accurate assessment of the Shannon diversity index is crucial, as microdiversity is determined as the exponent of the Shannon index (García-García et al., 2019). Each retained OTU was normalized to an overall abundance of 5000 reads for the calculation of microdiversity, persistence (i.e., the proportion of samples where an OTU/ASV was detected), and variance in abundance following methods described elsewhere (García-García et al., 2019; Linz et al., 2017). Principal Coordinates Analysis (PCoA) plot was used to visualize differences in OTU’s microdiversity in COM and RES samples.

### 2.4 Statistics analysis

Alpha-diversity (Shannon index) and beta-diversity (Bray-Curtis and Jaccard index) analyses were estimated using the phyloseq v. 1.46.0 (McMurdie and Holmes, 2013) and vegan v. 2.6.6.1 (Oksanen et al., 2001) packages. We employed distance-based redundancy analysis (dbRDA) to quantify the variance in the beta-diversity that could be explained by the measured environmental variables using the vegan package. Permutational multivariate analysis of variance (PERMANOVA) was performed using Adonis in vegan to compare the microbial community compositions in COM and RES samples. Mantel tests were used to estimate similarities between any two distance matrices (999 permutations, Spearman rank correlation). DESeq2 v. 1.42.1 (Love et al., 2014) was used on the unnormalized read count table of the retained OTUs to identify differentially abundant OTUs between COM and RES sampling sites. P-values were adjusted using the Benjamin and Hochberg method in DESeq2 (Benjamini and Hochberg, 1995). For all statistical analyses, the threshold of p < 0.05 was considered significant. All analyses were conducted in R v. 4.3.3 (R Core Team, 2013).

### 2.5 Community assembly mechanisms

The phylogenetic bin-based null model analysis via the iCAMP package (v. 1.5.12) (Ning et al., 2020) was used to quantitively infer the impact of stochastic and/or deterministic processes on ASV-based community assembly. This approach assigns the observed sequences into different phylogenetic bins based on a user-defined phylogenetic distance threshold of 0.2 (Ning et al., 2020). It then determines the assembly processes governing each phylogenetic bin based on null model analysis of the phylogenetic diversity using beta Net Relatedness Index (βNRI) and taxonomic β-diversities using modified Raup–Crick metric (RC). The assembly processes include homogeneous selection (HoS), heterogeneous selection (HeS), dispersal limitation (DL), homogenizing dispersal (HD), and drift (DR). Here, HD and DR are considered stochastic processes, and their combined relative importance was calculated as stochasticity. Pearson correlation was used to determine the relation between ecological processes and other variables following the methods described previously (Ning et al., 2024).

## 3. Results

### 3.1 Differences in water chemistry between RES and COM locations

A total of 100 drinking water samples were analyzed, including 60 COM samples and 40 RES samples from June to November 2020. Monthly water demand was 54,664 ± 25,205 m^3^/month in COM and 1,465 ± 457 m^3^/month in RES. The measured physical-chemical parameters are presented in Supporting Information (Table S2). RES samples generally demonstrated stable water pH (9.3 ± 0.1), DO (10.4 ± 0.6 mg/L), and total inorganic nitrogen (1.2 ± 0.2 mg N/L); these were similar to COM samples (9.3 ± 0.2, 10.8 ± 0.7 mg/L, and 1.2 ± 0.2 mg N/L). While total inorganic nitrogen had comparable concentrations in COM and RES samples, nitrite concentrations were consistently higher in RES (0.6 ± 0.09 mg N/L) than in COM samples (0.03 ± 0.03 mg N/L). Notable temporal changes were observed in the measured total chlorine concentration. In COM locations, the total chlorine concentration increased from 0.56 ± 0.69 mg/L in June to 2.1 ± 0.92 mg/L in October, then slightly decreased to 1.48 ± 0.89 mg/L in November. In RES locations, the total chlorine concentration increased from 1.6 ± 0.59 mg/L in June to 2.7 ± 0.03 mg/L in November. Overall, the RES samples presented markedly higher total chlorine concentration (2.3 ± 0.5 mg/L) than COM samples (1.3 ± 0.9 mg/L). Water temperature was also higher in COM (21 ± 2.4 °C) than in RES (19 ± 2.3 °C). The warmest month was August for RES locations (21.3 ± 0.3 °C) and June for COM locations (22.3 ± 2.3 °C).

### 3.2 Differences in microbial community compositions and factors influencing them

Drinking water microbial communities in RES and COM differed significantly (PERMANOVA, F = 16.8, R^2^ = 0.146, p = 0.001, 999 permutations). Based on the Jaccard index (Figure 1A), variance partitioning analysis revealed that the largest factor contributing to the overall beta-diversity was the building type (i.e., COM vs RES), explaining 10.5% of the total variance in beta-diversity (ANOVA, p = 0.001). Differences in individual buildings were also important, explaining 9.1% of the total variance (p = 0.001). The chlorine concentration was the most influential water chemistry parameter, which explained 4.5% of the total variance (p = 0.001). The sample collection month explained 4.2% of the total variance (p = 0.001). While these were the main factors, season (3.1% of total variance explained, p = 0.001), dissolved oxygen (1.7%, p = 0.003), pH (1.0%, p = 0.005), and temperature (1.0%, p = 0.006) all showed statistically significant effects on the observed variations. Variance partitioning analysis based on Bray-Curtis dissimilarity indicated similar patterns (Figure 1C). Building type (17.1%, p = 0.001), individual buildings (13.7%, p = 0.001), and chlorine concentrations (8.3%, p = 0.001) remained the most influential factors. Other factors mentioned above remained statistically significant to observed microbial variations. Pairwise beta-diversity for lumped COM and RES sample showed significant differences between the two building types when measured by the Jaccard index and Bray-Curtis dissimilarity (Figure 1B and 1D).

**Figure 1:**
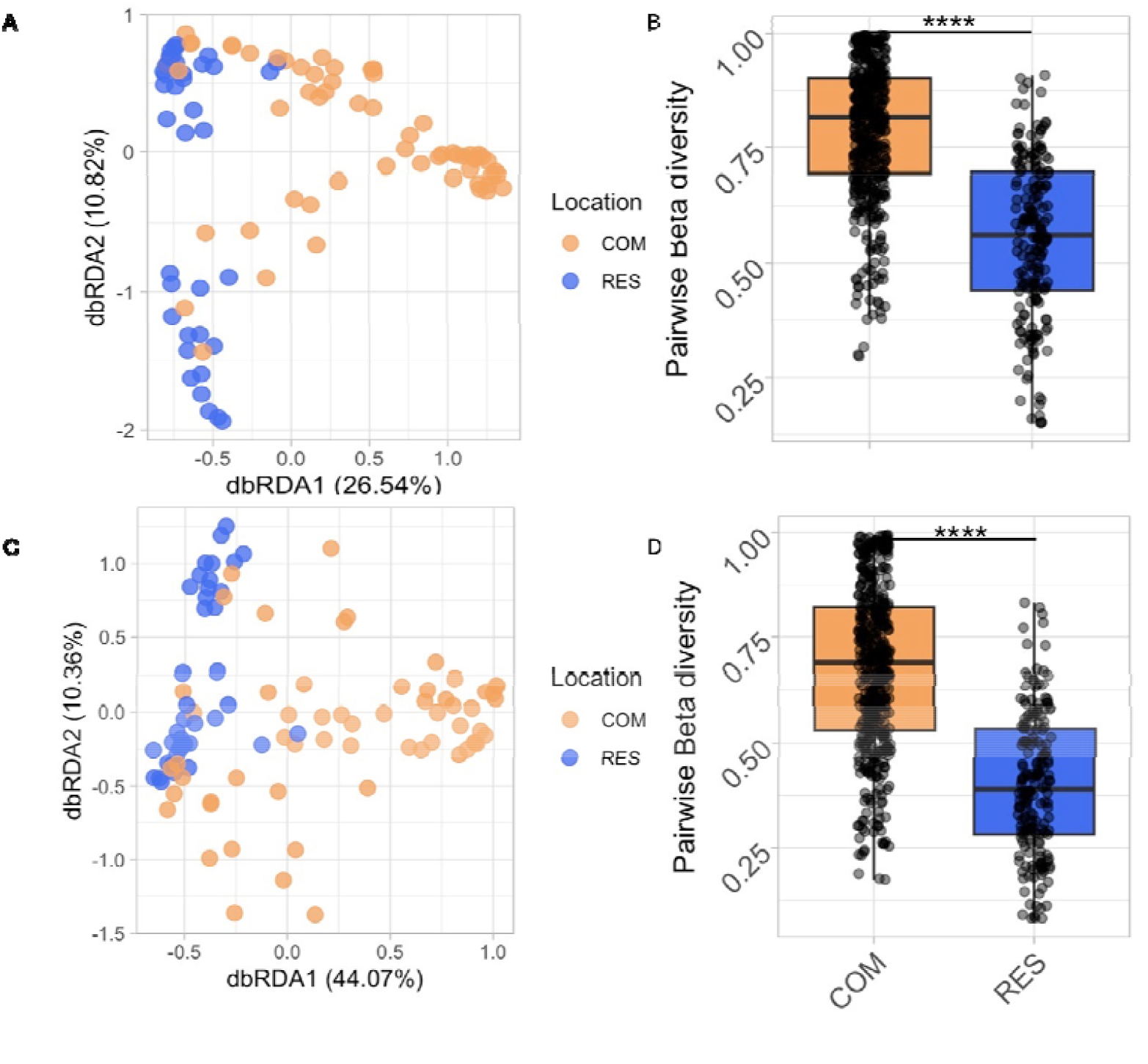
(A) Jaccard index-based distance-based redundancy analysis (dbRDA) plot for samples collected from different building types, COM: commercial buildings, RES: residential buildings. (B) Boxplots of the pairwise Jaccard index for lumped COM and RES sites. ****: T-Test p < 2.2e-16. (C) Bray-Curtis-based dbRDA plot for samples collected from different building types. (D) Boxplots of the pairwise Bray-Curtis dissimilarity for lumped COM and RES sites. ****: p < 2.2e-16.

### 3.3 Microdiversity, persistence, and variability

OTUs with total reads greater than 5000 were retained for further fine-scale microdiversity analysis. This step resulted in the retention of 49 OTUs across 100 samples. Despite the loss of sequences, the overall beta-diversity patterns were highly preserved (Figure S1), suggesting that these top 49 OTUs effectively captured the compositional shifts across the dataset. The COM and RES samples had distinct microdiversity profiles (Figure 2A) (PERMANOVA, F = 37.9, R^2^ = 0.279, p = 0.001). Interestingly, the Euclidean distance matrix constructed using the OTU microdiversity estimates was significantly correlated with the Bray-Curtis distance matrix based on the OTUs read abundance (Mantels r = 0.43, p = 0.001) (Figure 2B). This may suggest that the taxa driving the observed changes in overall beta-diversity were also driving the observed changes in microdiversity. In other words, the OTUs that were most responsible for the observed differences in community compositions were the same OTUs that exhibited the most significant within-species variation in microdiversity between COM and RES.

**Figure 2:**
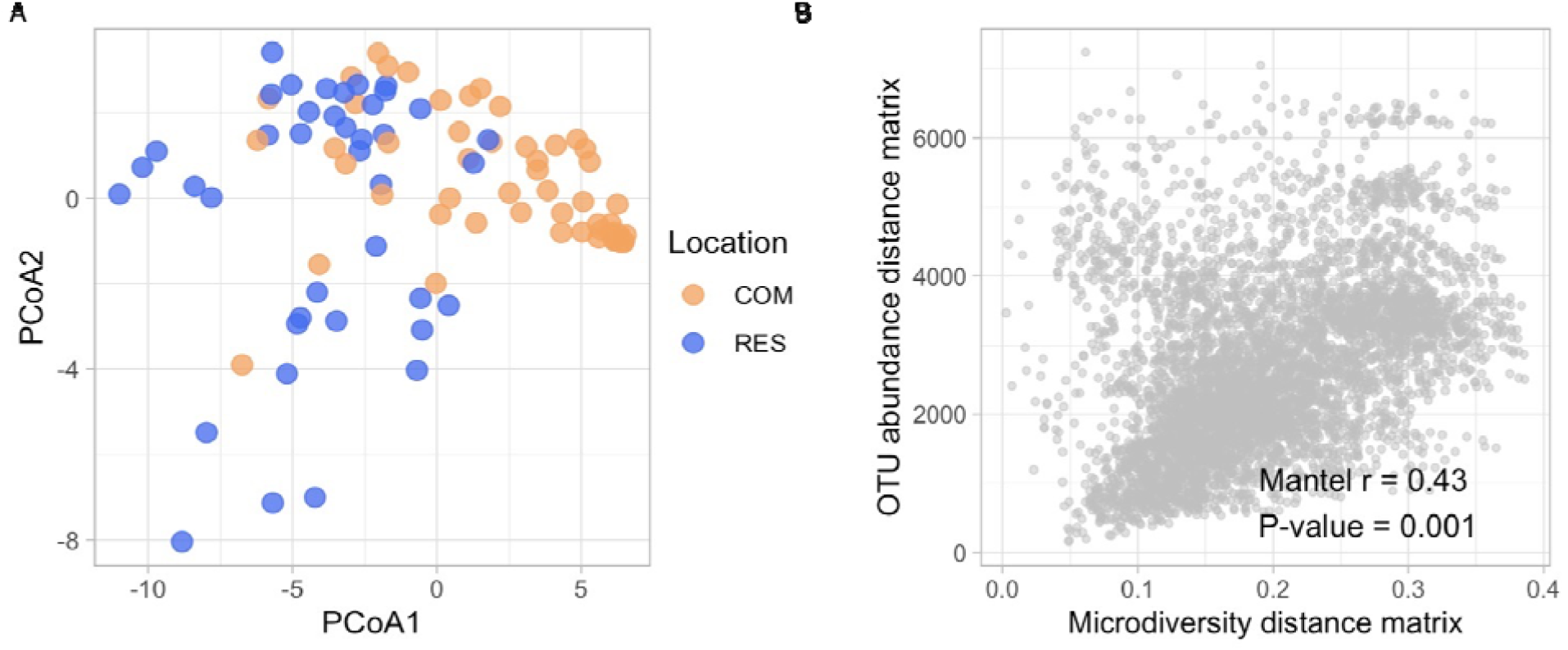
(A) Principal Coordinates Analysis (PCoA) plot based on the microdiversity of Operational Taxonomic Units (OTUs) in samples collected from different building types. Euclidean metrics were used to calculate dissimilarity. (B) Results of Mantel test between OTU read abundance matrix (Bray-Curtis) and microdiversity matrix (Euclidean). Mantel coefficient (r) and associated p-value were calculated.

OTUs exhibited significantly higher microdiversity in RES samples compared to COM sample (Figure 3A). ASV-rich OTUs were more frequently detected as compared to OTUs with fewer ASVs, exhibiting increased persistence (Figure 3B and 3E) and reduced variability in relative abundance (Figure 3C and 3F). A positive microdiversity-abundance relationship was observed in RES samples (Spearman r = 0.67, p = 1.5e-07) (Figure 3D). Conversely, this correlation wa not significant in COM samples (Spearman r = 0.096 p = 0.50) (Figure 3G).

**Figure 3:**
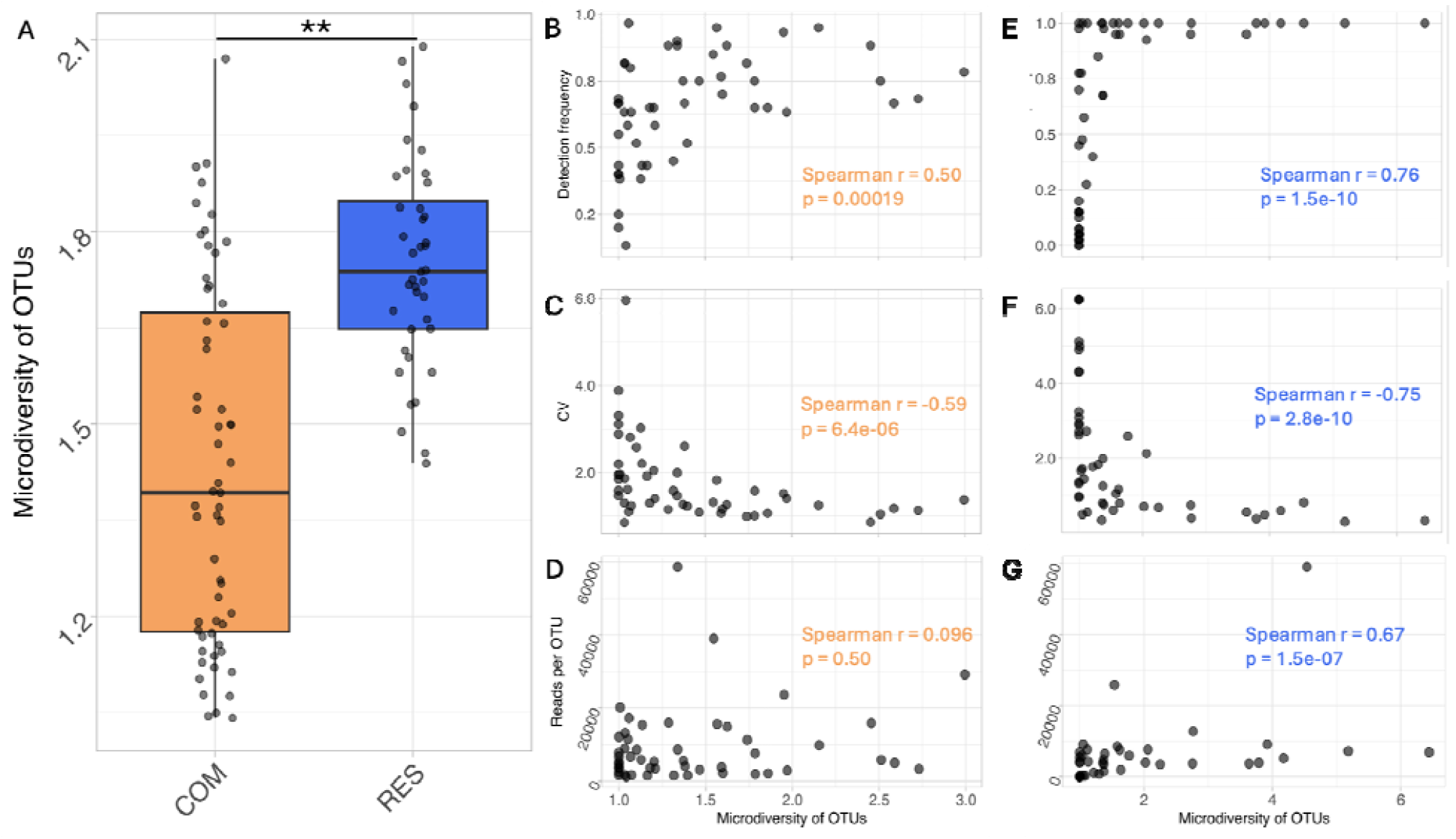
Mean effective microdiversity of Operational Taxonomic Units (OTUs) calculated across commercial (COM) and residential (RES) samples (**: p = 0.0099) (A). Scatter plots between microdiversity and OTU detection frequency (B and E), CV (C and F), and total reads per OTU (D and G). Spearman correlation strength r and p values are shown in the figure. Figure 3B, 3C, and 3D with yellow colored text are patterns within COM. Figures 3E, 3F, and 3G with blue-colored text are patterns within RES.

### 3.4 Microdiversity differences in differentially abundant OTUs in COM and RES samples

Of the 49 most abundant OTUs, DESeq2 identified 21 OTUs that were significantly more abundant in COM (Benjamin and Hochberg adjusted p < 0.05), 16 OTUs significantly more abundant in RES (p < 0.05), and 12 OTUs with no significant differences (p > 0.05) (Table S3). Figure 4 shows that OTUs found in higher abundance in samples from a specific building type (COM or RES) also showed greater microdiversity. It is important to note that microdiversity measures the *effective* ASVs rather than *constituent* ASVs, so the values were not integers. For OTUs that were significantly more abundant in RES, their microdiversity was 3.0 ± 1.6 in RES but only 1.7 ± 0.6 in COM (Figure 4A). Examples included *Duganella*-like (OTU 1683, 1838, 1867), *Undibacterium*-like (OTU 1667, 1840), and *Pseudomonas*-like OTUs (OTU 1478). OTUs that were significantly more abundant in COM had a microdiversity of 1.3 ± 0.3 in COM, whereas these OTUs contained only 1.0 ± 0.1 effective ASVs in RES samples (Figure 4B). Representatives included OTU 1979 (genus: *Curvibacter)*, OTU 2466 (genus: *Gemmatimonas)*,

**Figure 4:**
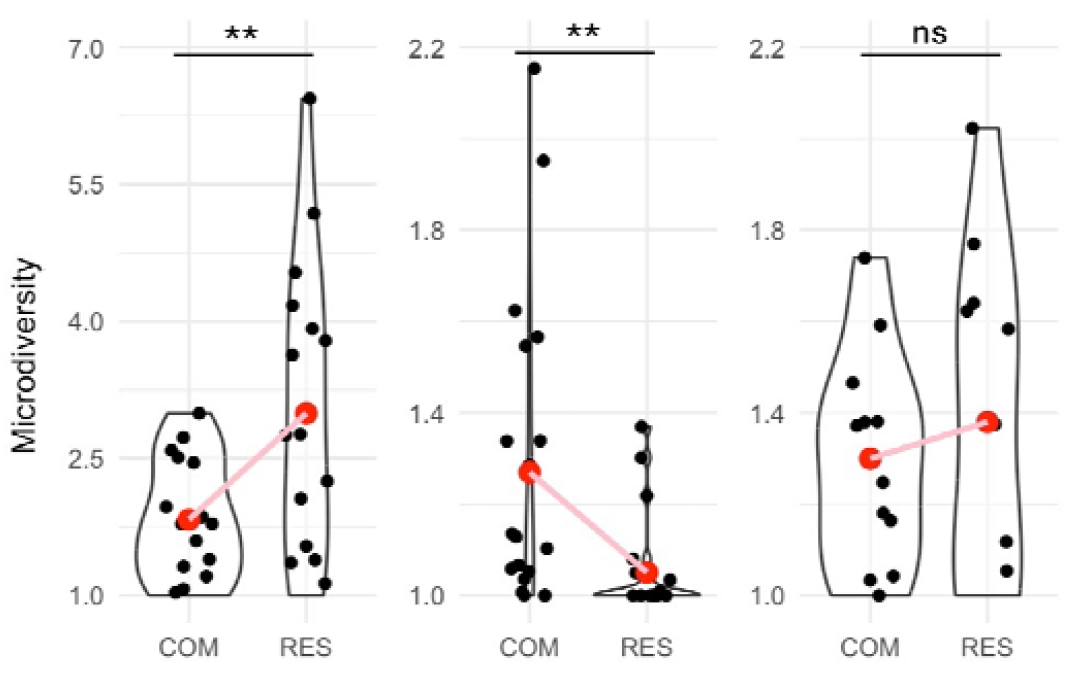
Microdiversity of three groups of Operational Taxonomic Units (OTUs) in commercial (COM) and residential (RES) samples. (A): the group of OTUs that were significantly more abundant in RES (n = 16), **: T-test, p = 0.0058. (B): OTUs significantly more abundant in COM (n = 21), **: p = 0.0033. (C): OTUs that did not show significant differences in abundances between building types (n = 12), ns: no significance (p = 0.15).

### 3.5 Spatial distributions and temporal dynamics of ASVs within the same OTU

ASVs within the same OTU showed distinct spatial distributions (Figure 5). For instance, for OTU 1874 (species: *Acidovorax delafieldii*), OTU 1992 (genus: *Sediminibacterium*), OTU 2466 (genus: *Gemmatimonas*), and OTU 4423 (genus: *Sphingomonas*), their most abundant ASV differed in COM and RES. Further, the number of ASVs per OTU also varied spatially. For OTU 1874, while four different ASVs were detected in COM with ASV 9 as the highest abundant (and persistent) sub-taxa, ASV 632 was the only sub-taxa within OTU 1874 in RES locations. In addition, while ASV 9 was present in 83% of the COM samples, ASV 632 appeared in only half of the RES samples.

**Figure 5:**
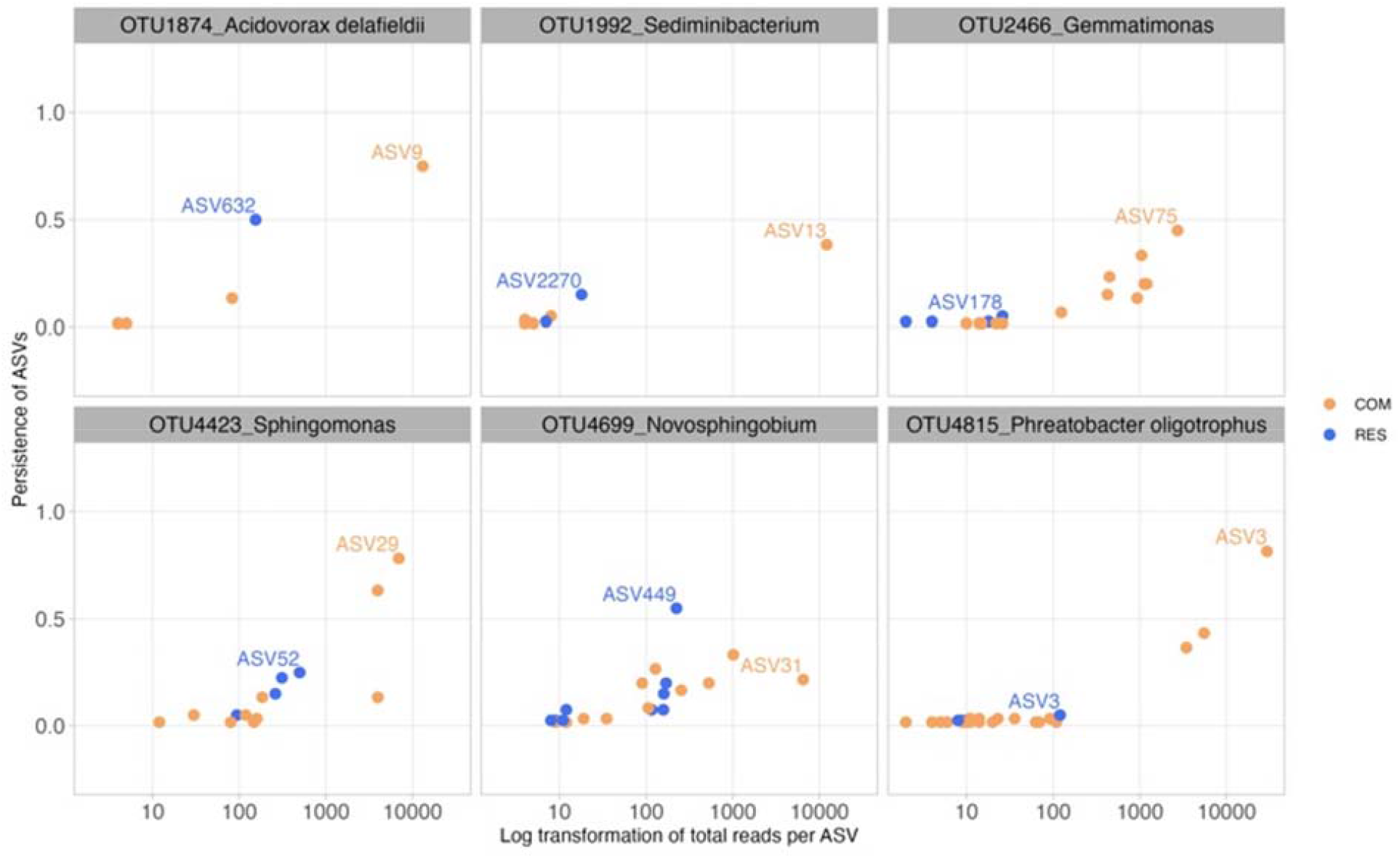
Scatter plots of log-transformed global abundance and persistence for constituent Amplicon Sequence Variants (ASVs) of each Operational Taxonomic Unit (OTU) in commercial (COM, yellow) and residential (RES, blue) samples. Each dot represents a single ASV, and the most abundant ASVs in COM and RES samples were labeled with ASV-IDs respectively.

ASVs within the same OTU exhibited varied temporal patterns. Figure 6 shows the decomposition of OTU 1486 (most abundant, family: Neisseriaceae), OTU 1492 (most microdiverse, genus: Rhodoferax), and OTU 1683 (most divergent in microdiversity between building types, genus *Duganella*) into main constituent ASVs, whose accumulated abundance accounted for at least 50% of the OTU’s abundance. ASV 2 was the most dominant and abundant sub-taxa of OTU 1486 in both building types from June to July, accounting for 45 ± 15 % of OTU 1486’s abundance in COM and 42 ± 12% in RES (Figure 6A, 6B). However, ASV 19 became more abundant in August and was the dominant sub-taxa of OTU1484 in later months (COM: 37 ± 12% and RES: 33 ± 12%). For OTU 1492 (Figure 6C, 6D), ASV 145 was present in both types of buildings from June to August but was no longer detectable in the following months. In contrast, ASV 124 of OTU 1492 persisted throughout the entire period in both type of buildings. In the case of OTU 1683, ASV 294 and ASV 305 both presented in COM from July to October, but ASV 294 was not detected in June and ASV 305 was not detected in November (Figure 6E). A higher degree of microdiversification of OTU 1683 was seen in RES (Figure 6F). Notably, ASV 326 was only present in June to August and was not present in later months.

**Figure 6:**
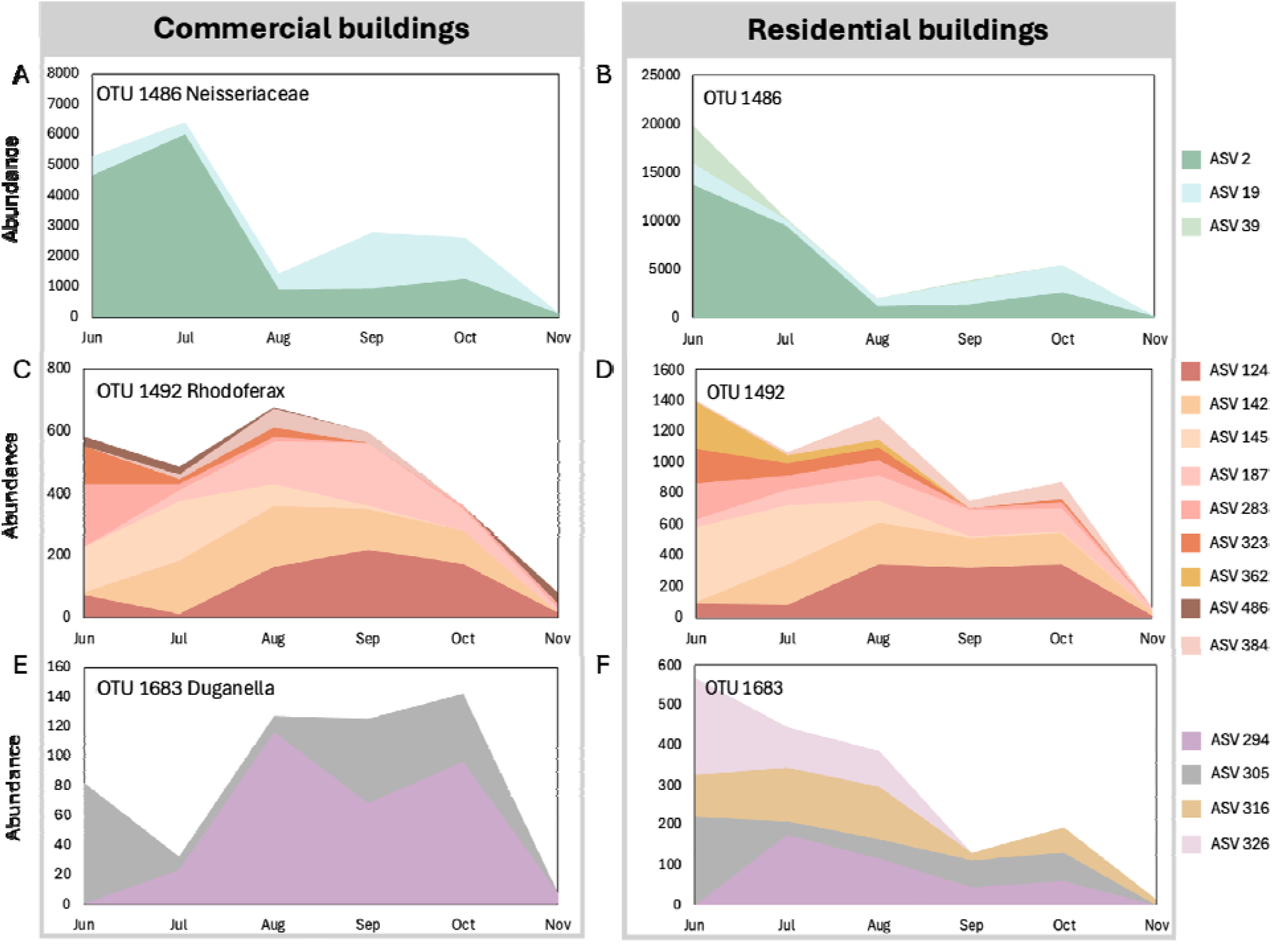
Temporal change of constituent ASVs of OTU 1486 (A and B), OTU 1492 (C and D), and OTU 1683 (E and F) in commercial and residential buildings. ASV: Amplicon Sequence Variant. OTU: Operational Taxonomic Unit.

### 3.6 Ecological forces shaping building plumbing associated communities

The distinct community compositions and microdiversity in COM and RES prompted further inquiry into the ecological forces driving the observed patterns. Dispersal limitation (DL) and drift (DR) were dominant ecological processes in COM communities throughout the entire study period, with the relative importance of 25.0-54.5% and 28.3-71.6%, respectively (Figure 7A). Among the eight phylogenetic bins (Table S4) clustered from 438 ASVs (Figure 7C), three bins were largely driven by DL. Their top taxa were affiliated to *Duganella* (bin1, relative importance of DL= 68.1%), *Rugamonas* (bin2, 44.6%), and *Paucibacter* (bin8, 47.2%), respectively. From June to November, we generally observed increasing stochasticity in COM communities, with the relative importance rising from 50.1% in June to 72.0% in November.

**Figure 7:**
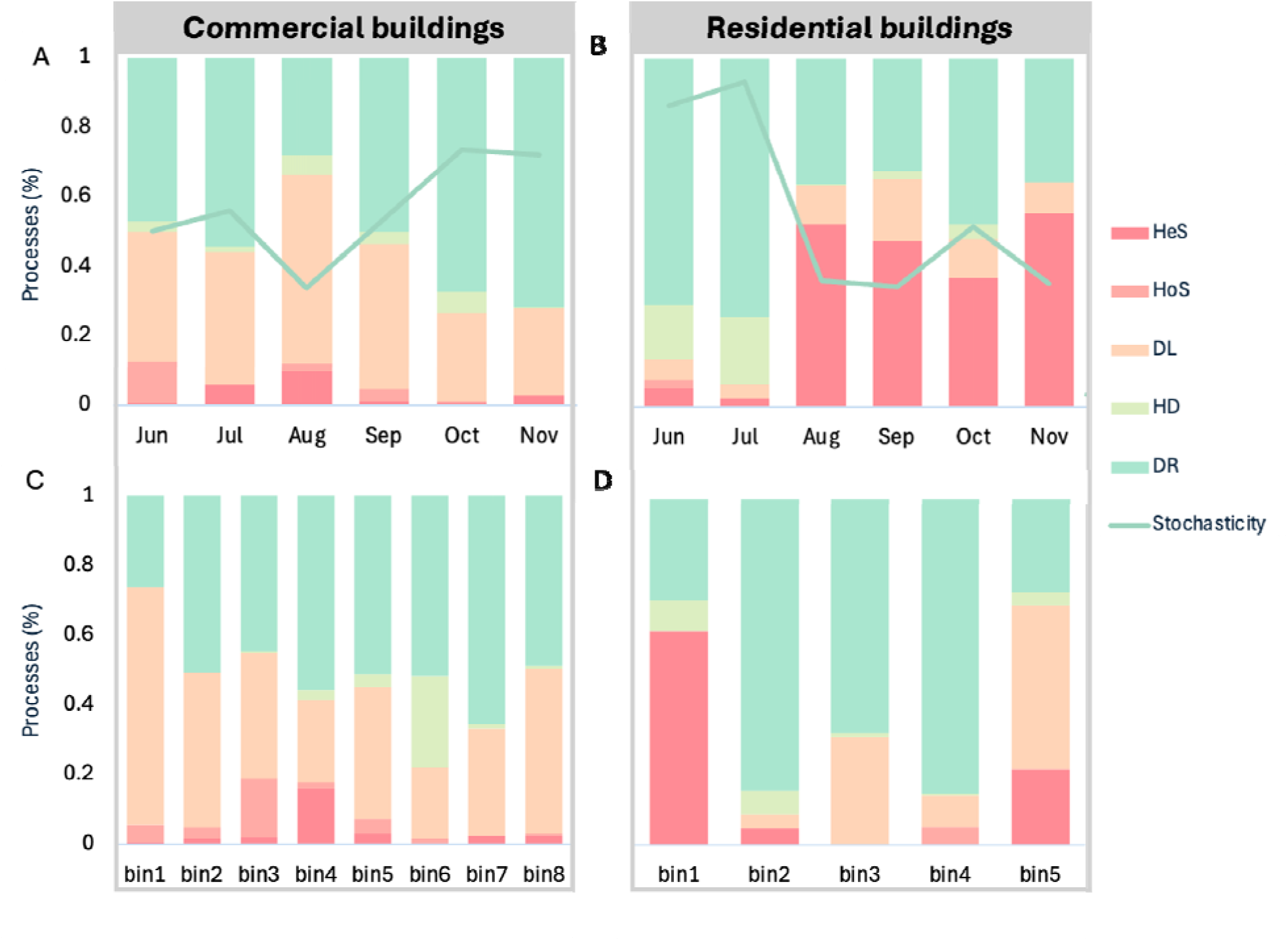
Assembly processes in commercial and residential tap water communities in each month (A and B) and different phylogenetic groups (C and D). HeS: heterogeneous selection; HoS: homogeneou selection; DL: dispersal limitation; HD: homogeneous dispersal; DR: drift.

In RES samples, the relative importance of stochasticity dropped from 86.3% in June to 35.4% in November (Figure 7B). 337 ASVs were clustered into five phylogenetic bins (Figure 7D, Table S5). DR was found to be the most important ecological process in June and July (relative important = 70.8-74.2%) and was the dominant force influencing three bins. Deterministic processes, mainly HeS (36.9-55.8%), played a more dominant role from August to November. Bin 1 with the top taxon affiliated to *Duganella paludism* was the most influential phylogenetic group driving the community assembly in RES samples. It had a relative abundance of 57.4% among all taxa in RES samples and contributed 43.9% to the observed temporal shift from DR to HeS as the primary community-shaping mechanism. DL also accounted for an increased proportion from August to November (8.8-17.8%) (Figure 7B), and DL was consistently the dominant force in bin 5 affiliated with *Mycobacterium frederiksbergense* during this time. Detailed information regarding taxa, bin ID, contribution proportions, and dominant processe are in Table S4-S5. Pearson correlation revealed that in COM samples, pH was significantly correlated with DL (r = -0.94, p = 0.016), and total nitrogen (r = -0.49, p = 0.025) wa significantly associated with DR. Interestingly, in RES samples, while several environmental variables showed a strong correlation with the observed assembly pattern, such as dissolved oxygen and total chlorine concentrations, none of the correlations was statistically significant (p > 0.05). Detailed data are presented in Figure S2.

## 4. Discussion

### 4.1 Distinct environments in plumbing systems lead to building-type-specific microbial communities

Throughout the study period, COM samples had a lower chlorine residual and warmer temperatures compared to RES samples. This is common in plumbing systems of large buildings and is associated biofilm development and microbial regrowth (Bédard et al., 2016; Greenwald et al., 2022; Hozalski et al., 2020). Our previous study reported a gradual convergence in COM and RES microbial communities post-reopening after extended COM building closures during the COVID-19 pandemic, indicating that the resumption of water use played a key role in the recovery of building plumbing-associated microbial communities (Vosloo et al., 2023). Despite this convergence, COM and RES microbial communities remained distinct during the entire sampling period (Figure 1A and 1C). Among individual sites within the same building type, the microbial communities in COM building plumbing showed greater heterogeneity compared to those in RES buildings. The trend was consistent with both measurements considering taxa abundance (Bray–Curtis) (Figure 1B) and sole presence/absence of taxa (Jaccard index) (Figure 1D). The higher beta diversity in COM buildings likely emerges not only from the greater complexity of the building plumbing compared to RES but also due to the presence of diverse microbial niches, resulting in more unique community membership and differential abundance of taxa shared with RES drinking water samples.

### 4.2 Microdiversity is associated with the preservation of the higher-order community in building plumbing

Microdiversity may play an important role in preserving the higher-order community in building plumbing. For instance, different ASVs within the same OTU exhibited different habitat preferences (i.e., COM vs RES) (Figure 5) while also exhibiting building-specific temporal variability (Figure 6). However, regardless of the spatial-temporal ASV turnover, the higher taxonomy groups (OTUs) with more ASVs exhibited higher persistence. For any taxa, the presence of multiple sub-taxa can decrease the likelihood that the strain-specific disadvantage of a single sub-taxa results in the decline of the whole population (Needham et al., 2017). OTUs with a greater microdiversity demonstrated higher persistence and lower variability in relative abundance compared to OTUs with fewer ASVs (Figure 3). Similar trends have been observed in freshwater (García-García et al., 2019; Linz et al., 2017; Okazaki et al., 2021) and marine taxa (Mende et al., 2019; Needham et al., 2017). Further, we also find that OTUs with a larger number of ASVs also tend to exhibit higher overall relative abundance (Figure 4A and 4B).

These observations are consistent with the theory that microdiversity enhances the overall species fitness (Acinas et al., 2004). One potential explanation for OTUs with reduced microdiversity could be a genetic sweep (Acinas et al., 2004; Needham et al., 2017), whereby the evolution of specific adaptive traits within a strain results in a substantial fitness differential between strains and thus, the exclusion of the other strains (Needham et al., 2017). Further, the existence of niche selection among strains from the same species has been documented in a broad range of ecosystems, including soil (Larkin and Martiny, 2017), glacier-fed streams (Fodelianakis et al., 2022), freshwater (García-García et al., 2019; Linz et al., 2017; Okazaki et al., 2021), fermented milk (You et al., 2023), marine (Needham et al., 2017), and activated sludge (Cotto et al., 2023; Srinivasan et al., 2021). The unique spatial and temporal dynamics observed among ASVs within the same OTU, both in this study and in others, indicate that this phenomenon can be generalized to the drinking water microbiome.

### 4.3 Microbial communities within the plumbing system displayed building type-specific assembly patterns

Dispersal limitation was the dominant ecological mechanism shaping COM drinking water microbial communities compared to heterogeneous selection in RES samples (Figures 7A and 7B). The importance of dispersal limitation in building plumbing microbiomes has been reported before (Cai et al., 2023; Liu et al., 2024), and we further revealed that its impact can be higher or lower depending on the building type. The strong role of dispersal limitation in COM suggests these microbial communities are highly localized and largely isolated with microbial compositions less impacted by immigration from the DWDS over sampling time scales (Ling et al., 2018). Dispersal limitation primarily governed the turnover in three phylogenetic bins (Figure 7C). On a broader taxonomy scale, members within these bins belong to the family *Oxalobacteraceae* and *Comamonadaceae*; both bacteria from families have been implicated in biofilm formation (Douterelo et al., 2014b; Nguyen et al., 2023). We hypothesize that local biofilm seeding and regrowth due to stagnation contribute to shaping the bulk water community in COM plumbing. Specific to this study, the extended water stagnation and disruption in routine water usage patterns in COM due to the COVID-19 pandemic may have further exacerbated the impact of dispersal limitation. These factors may explain the observed lower microdiversity in COM samples (Figure 3A) relative to RES samples. Biofilm seeding and water stagnation can reduce microbial diversity due to the selection of specific taxa (Douterelo et al., 2016, 2014a; Proctor et al., 2018). These insights on building type-specific community assembly forces could help with tailoring building water management practices. For instance, two similarly sized COM buildings may require markedly different flushing durations based on the extent to which their building plumbing communities are shaped by dispersal limitation.

Microbial community dynamics in RES samples indicated a high influence of heterogeneous selection (Figure 7B); i.e., selective forces that shape microbial communities towards different community structures over spatial-temporal gradients. Selection pressures are deterministic and can often be related to definable variables (Zhou and Ning, 2017). However, we found no evident impact of the measured environmental parameters on the heterogeneous selection governing the residential communities (Figure S2). This may suggest that the microbial communities in RES plumbing were less influenced by the observed moderate changes in disinfectant stress or substrate availability in bulk water. Drinking water treatment (e.g., filtration, disinfection) imposes significant selective pressures on the drinking water microbiome (El-Chakhtoura et al., 2015; Pinto et al., 2012; Siponen et al., 2024; Thom et al., 2022; Zheng et al., 2024). As a result, the small variation in water chemistry parameters between RES building plumbing may not have as strong of an impact on shaping the RES microbial communities (Guo et al., 2019; Richter-Heitmann et al., 2020). Because heterogeneous selection (i.e., a deterministic mechanism) was the primary driver of community assembly in RES locations, and none of the measured environmental parameters are correlated with it, this raises the possibility that other unmeasured factors are likely important. Indeed, factors such as plumbing design (Ning et al., 2024; Pinto et al., 2014; Stegen et al., 2013) as well as properties like age, diameter, and materials (Cruz et al., 2020; Lee et al., 2021; Ling et al., 2018; Schück et al., 2023) might be primary drivers of community assembly in RES locations.

## 5. Conclusions

We used full-length 16S rRNA gene sequencing to systematically compare microbial communities in RES and COM building plumbing. In doing so, we identified critical insights on the impact of microdiversity on population persistence and abundance, as well as the differing roles of ecological mechanisms shaping building plumbing microbial communities.

- Microdiversity of the drinking water microbiome is strongly influenced by building type.
- Higher microdiversity is associated with higher persistence and relative abundance
- ASVs within an OTU demonstrate habitat preferences based on building types
- Different ecological mechanisms shape COM and RES building plumbing microbial communities

## Acknowledgment

This research is supported by National Science Foundation Grants CBET 2029850, 2220792, and USEPA Grant R840606.

## Data Availability

The raw FCS files have been deposited in the Flow Repository and are publicly available under accession number FR-FCM-Z4SR, and 16S rRNA gene sequences are available on NCBI at BioProject accession number PRJNA885853. All code for amplicon sequencing, microdiversity analysis, statistical analysis, and plots are available upon request.

