## Supplemental Material for "Building plumbing influences the microdiversity and community assembly of the drinking water microbiome"

**Table S1: Summary of reads per sample at different stages of data processing with DADA2 (Control = blank filters, SW = sampled water)**

| **Sample ID** | **After circular Control sensus sequencing** | **Post primer removal** | **Post quality filtering** | **Post denoising** |
| --- | --- | --- | --- | --- |
| CONTROL 01 | 48 | 41 | 38 | 4 |
| CONTROL 02 | 34 | 29 | 27 | 3 |
| CONTROL 03 | 55 | 50 | 46 | 8 |
| CONTROL 04 | 14 | 14 | 8 | 2 |
| CONTROL 05 | 356 | 315 | 311 | 249 |
| CONTROL 06 | 18 | 15 | 12 | 3 |
| CONTROL 07 | 17 | 16 | 14 | 1 |
| CONTROL 08 | 29 | 28 | 26 | 9 |
| CONTROL 09 | 33 | 29 | 24 | 6 |
| CONTROL 10 | 23 | 19 | 17 | 1 |
| CONTROL 11 | 22 | 21 | 17 | 1 |
| CONTROL 12 | 25 | 21 | 21 | 2 |
| CONTROL 13 | 34 | 29 | 28 | 1 |
| CONTROL 14 | 21 | 17 | 16 | 1 |
| CONTROL 15 | 16 | 12 | 9 | 1 |
| CONTROL 16 | 35 | 32 | 30 | 3 |
| CONTROL 17 | 26 | 22 | 21 | 1 |
| CONTROL 18 | 27 | 26 | 24 | 2 |
| CONTROL 19 | 19 | 18 | 17 | 1 |
| CONTROL 20 | 10 | 10 | 7 | 1 |
| CONTROL 21 | 25 | 24 | 20 | 9 |
| CONTROL 22 | 10 | 9 | 8 | 1 |
| CONTROL 23 | 6 | 4 | 3 | 1 |
| CONTROL 24 | 18 | 14 | 12 | 4 |
| SW001 | 26753 | 23226 | 22880 | 18022 |
| SW002 | 22718 | 19594 | 19244 | 13874 |
| SW003 | 29465 | 25182 | 24784 | 20636 |
| SW004 | 24937 | 21734 | 21413 | 17739 |
| SW007 | 32342 | 27966 | 27530 | 22889 |
| SW008 | 29367 | 25504 | 25092 | 20278 |
| SW009 | 33634 | 28836 | 28464 | 24059 |
| SW010 | 27430 | 23345 | 22940 | 17998 |
| SW011 | 31229 | 27042 | 26754 | 24247 |
| SW012 | 20486 | 17999 | 17836 | 16356 |
| SW013 | 4145 | 3672 | 3645 | 3568 |
| SW014 | 28005 | 24660 | 24284 | 19523 |
| SW015 | 32825 | 28436 | 28075 | 24782 |
| SW016 | 34492 | 30042 | 29596 | 25221 |
| SW024 | 19184 | 16491 | 16311 | 14943 |
| SW025 | 19014 | 16622 | 16451 | 13841 |
| SW026 | 21870 | 19541 | 19299 | 18493 |
| SW027 | 32103 | 27800 | 26601 | 23873 |
| SW028 | 20797 | 18127 | 17913 | 16394 |
| SW029 | 24273 | 20778 | 20467 | 16311 |
| SW030 | 24323 | 21110 | 20786 | 15967 |
| SW031 | 24658 | 21489 | 21135 | 16070 |
| SW032 | 25246 | 22317 | 22012 | 18940 |
| SW033 | 23870 | 20486 | 20238 | 16960 |
| SW036 | 45 | 42 | 42 | 8 |
| SW037 | 27241 | 23721 | 23367 | 19192 |
| SW038 | 30061 | 26510 | 26099 | 21101 |
| SW039 | 33938 | 29249 | 28749 | 23013 |
| SW040 | 34930 | 30013 | 29624 | 25442 |
| SW041 | 38848 | 34111 | 33735 | 30213 |
| SW042 | 28615 | 24532 | 24243 | 22244 |
| SW043 | 32207 | 28398 | 28000 | 22805 |
| SW044 | 33131 | 27379 | 27057 | 24738 |
| SW045 | 34550 | 30153 | 29649 | 23906 |
| SW053 | 470 | 417 | 413 | 270 |
| SW054 | 33124 | 29134 | 28742 | 23318 |
| SW055 | 21829 | 19221 | 19047 | 16176 |
| SW056 | 28927 | 25816 | 25441 | 20253 |
| SW057 | 28261 | 24869 | 24479 | 23583 |
| SW058 | 39341 | 34752 | 32943 | 28167 |
| SW059 | 32442 | 27812 | 27326 | 20164 |
| SW060 | 32248 | 28023 | 27504 | 20120 |
| SW061 | 26686 | 23293 | 23000 | 18977 |
| SW062 | 23791 | 21030 | 20740 | 16833 |
| SW065 | 30265 | 26518 | 26157 | 20826 |
| SW066 | 34193 | 29657 | 29154 | 21666 |
| SW067 | 28278 | 24801 | 24387 | 19332 |
| SW068 | 31358 | 27730 | 27256 | 21394 |
| SW069 | 22527 | 19622 | 19407 | 16496 |
| SW070 | 25731 | 22502 | 22267 | 18667 |
| SW071 | 38579 | 32834 | 32397 | 30432 |
| SW072 | 33821 | 29283 | 28857 | 22659 |
| SW073 | 30002 | 25607 | 25278 | 22209 |
| SW074 | 34498 | 29875 | 29371 | 22507 |
| SW082 | 22181 | 19387 | 19170 | 14822 |
| SW083 | 25630 | 22210 | 21874 | 16048 |
| SW084 | 36065 | 31207 | 30607 | 27659 |
| SW085 | 34620 | 29879 | 29434 | 23582 |
| SW086 | 400 | 352 | 349 | 245 |
| SW087 | 30008 | 26074 | 25682 | 19398 |
| SW088 | 24493 | 21020 | 20710 | 14842 |
| SW089 | 23948 | 20901 | 20625 | 15942 |
| SW090 | 18214 | 15756 | 15589 | 12616 |
| SW091 | 14175 | 12760 | 12598 | 10196 |
| SW094 | 20364 | 17597 | 17402 | 14178 |
| SW095 | 20208 | 17560 | 17338 | 13757 |
| SW096 | 23586 | 20350 | 20101 | 16232 |
| SW097 | 25842 | 22526 | 22183 | 16853 |
| SW098 | 33980 | 28642 | 28261 | 25275 |
| SW099 | 30397 | 26623 | 26239 | 20186 |
| SW100 | 17495 | 15281 | 15130 | 12565 |
| SW101 | 21404 | 19231 | 19013 | 15339 |
| SW102 | 29480 | 25489 | 25192 | 22208 |
| SW103 | 20552 | 17779 | 17585 | 14218 |
| SW111 | 8390 | 7302 | 7242 | 5732 |
| SW112 | 21458 | 18652 | 18427 | 14504 |
| SW113 | 34314 | 30823 | 29509 | 27914 |
| SW114 | 17659 | 15203 | 14992 | 13095 |
| SW115 | 8906 | 7911 | 7834 | 6224 |
| SW116 | 22867 | 19766 | 19519 | 15478 |
| SW117 | 27792 | 24428 | 24059 | 18674 |
| SW118 | 20828 | 18515 | 18234 | 13589 |
| SW119 | 21370 | 19030 | 18766 | 14731 |
| SW120 | 24889 | 21840 | 21480 | 16669 |
| SW123 | 27791 | 24944 | 24617 | 19650 |
| SW124 | 19157 | 16930 | 16727 | 13074 |
| SW125 | 25385 | 22754 | 22494 | 18287 |
| SW126 | 20043 | 17561 | 17307 | 13762 |
| SW127 | 46723 | 39538 | 38938 | 35136 |
| SW128 | 31303 | 27444 | 27019 | 20776 |
| SW129 | 9869 | 8595 | 8517 | 6426 |
| SW130 | 22279 | 19891 | 19669 | 15667 |
| SW131 | 37091 | 31230 | 30741 | 26222 |
| SW132 | 29983 | 26479 | 26049 | 19061 |
| SW140 | 14993 | 13221 | 13092 | 10602 |
| SW141 | 27323 | 23931 | 23583 | 16754 |
| SW142 | 15123 | 12468 | 12355 | 11900 |
| SW143 | 23666 | 21322 | 20955 | 15918 |
| SW144 | 13580 | 12105 | 11996 | 9659 |
| SW145 | 22245 | 19835 | 19559 | 15166 |
| SW146 | 10543 | 9284 | 9205 | 7365 |
| SW147 | 15478 | 13789 | 13635 | 11148 |
| SW148 | 15970 | 14236 | 14056 | 11525 |
| SW149 | 12777 | 11418 | 11263 | 9342 |
| SW152 | 19553 | 17351 | 17167 | 14591 |
| SW153 | 16465 | 14433 | 14274 | 11534 |
| SW154 | 17093 | 15259 | 15073 | 12415 |
| SW155 | 15324 | 13565 | 13377 | 11019 |
| SW156 | 30754 | 26647 | 26343 | 24781 |
| SW157 | 22432 | 19457 | 19257 | 16637 |
| SW158 | 16695 | 15192 | 15059 | 13427 |
| SW159 | 23246 | 20644 | 20437 | 17999 |
| SW160 | 39588 | 34752 | 34282 | 31086 |
| SW161 | 32590 | 28493 | 28038 | 25128 |
| SW169 | 1623 | 1411 | 1392 | 984 |
| SW170 | 21513 | 18994 | 18778 | 15735 |
| SW171 | 26016 | 23038 | 22617 | 21455 |
| SW172 | 17064 | 15103 | 14934 | 12933 |
| SW173 | 2048 | 1827 | 1814 | 1404 |
| SW174 | 13276 | 11748 | 11627 | 9517 |

**Table S2: Sample information including physiochemical parameters, location, collection month, and monthly water demand associated with buildings.**

| Sample | Property | Building type | Season | Temperature  (°C) | pH | Conductivity  (μS/m) | Dissolved oxygen  (mg/L) | Total chlorine  (mg/L) | Ammonium  (mg-N/L) |
| --- | --- | --- | --- | --- | --- | --- | --- | --- | --- |
| SW001 | RES01 | RES | Summer | 22.3 | 9.3 | 235.2 | 11.54 | 1.86 | 0.46 |
| SW002 | RES01 | RES | Summer | 17 | 9.3 | 228 | 11.07 | 2.16 | 0.48 |
| SW003 | RES02 | RES | Summer | 19.3 | 9.3 | 228.7 | 10.95 | 2.16 | 0.51 |
| SW004 | RES02 | RES | Summer | 16 | 9.2 | 229.2 | 10.42 | 2.24 | 0.51 |
| SW007 | RES04 | RES | Summer | 23.8 | 9.3 | 229.8 | 12.04 | 0.93 | 0.46 |
| SW008 | RES04 | RES | Summer | 15.5 | 9.3 | 228.5 | 12.57 | 1.12 | 0.47 |
| SW009 | RES05 | RES | Summer | 23.9 | 9.3 | 228.8 | 11.72 | 0.93 | 0.45 |
| SW010 | RES05 | RES | Summer | 14.9 | 9.2 | 229 | 12.4 | 1.13 | 0.47 |
| SW011 | COM01 | COM | Summer | 24 | 9.1 | 229.3 | 12.56 | 0 | 0.47 |
| SW012 | COM01 | COM | Summer | 23.3 | 9.2 | 229 | 13.03 | 0.15 | 0.45 |
| SW014 | COM02 | COM | Summer | 18.5 | 9.3 | 228.1 | 12 | 0.88 | 0.45 |
| SW015 | COM03 | COM | Summer | 26.7 | 8.6 | 234.7 | 12.25 | 0.15 | 0.37 |
| SW016 | COM03 | COM | Summer | 23 | 9.3 | 237.2 | 14.77 | 1.35 | 0.39 |
| SW024 | COM04 | COM | Summer | 24.9 | 9.1 | 233.9 | 12.47 | 0.08 | 0.49 |
| SW025 | COM04 | COM | Summer | 27.3 | 9.2 | 230.4 | 13.31 | 0.4 | 0.43 |
| SW026 | COM05 | COM | Summer | 20.9 | 8.7 | 226.8 | 8.23 | 0.01 | 0.41 |
| SW027 | COM05 | COM | Summer | 20.4 | 9.3 | 228 | 11.75 | 1.02 | 0.43 |
| SW028 | COM06 | COM | Summer | 20.1 | 9.3 | 232.5 | 10.71 | 0.02 | 0.48 |
| SW029 | COM06 | COM | Summer | 16.1 | 9.4 | 228.8 | 11.64 | 2.08 | 0.47 |
| SW030 | RES01 | RES | Summer | 23.9 | 9.3 | 192.6 | 9.97 | 1.61 | 0.45 |
| SW031 | RES01 | RES | Summer | 20.6 | 9.4 | 191.1 | 9.81 | 2.32 | 0.48 |
| SW032 | RES02 | RES | Summer | 24.2 | 9.7 | 190.3 | 9.72 | 2.32 | 0.5 |
| SW033 | RES02 | RES | Summer | 17.6 | 9.3 | 186.3 | 9.97 | 2.46 | 0.5 |
| SW037 | RES04 | RES | Summer | 17.5 | 9.4 | 187.4 | 11.36 | 2.26 | 0.5 |
| SW038 | RES05 | RES | Summer | 26.2 | 9.4 | 190.3 | 10.78 | 2 | 0.46 |
| SW039 | RES05 | RES | Summer | 17.2 | 9.4 | 188 | 11.13 | 2.36 | 0.49 |
| SW040 | COM01 | COM | Summer | 24.6 | 9.3 | 195.2 | 11.88 | 0.03 | 0.52 |
| SW041 | COM01 | COM | Summer | 25.6 | 9.2 | 195.2 | 11.85 | 0.02 | 0.51 |
| SW042 | COM02 | COM | Summer | 24.4 | 9.1 | 200.6 | 10.98 | 0.5 | 0.52 |
| SW043 | COM02 | COM | Summer | 21.6 | 9.4 | 191.2 | 11.26 | 1.84 | 0.48 |
| SW044 | COM03 | COM | Summer | 21.4 | 9.2 | 188.1 | 11.74 | 1 | 0.6 |
| SW045 | COM03 | COM | Summer | 21.9 | 9.4 | 179.3 | 12.51 | 1.73 | 0.61 |
| SW054 | COM04 | COM | Summer | 27 | 9.4 | 181.3 | 10.41 | 1.54 | 0.66 |
| SW055 | COM05 | COM | Summer | 21.1 | 8.9 | 207.3 | 9.33 | 0.02 | 0.33 |
| SW056 | COM05 | COM | Summer | 18.7 | 9.5 | 177.1 | 10.52 | 1.95 | 0.67 |
| SW057 | COM06 | COM | Summer | 20.9 | 9.9 | 181 | 9.85 | 0.81 | 0.63 |
| SW058 | COM06 | COM | Summer | 17.2 | 9.6 | 177.5 | 10.79 | 2.28 | 0.3 |
| SW059 | RES01 | RES | Summer | 21.5 | 9.3 | 214.4 | 9.31 | 1.74 | 0.58 |
| SW060 | RES01 | RES | Summer | 21.6 | 9.3 | 213.5 | 9.47 | 2.44 | 0.57 |
| SW061 | RES02 | RES | Summer | 21.1 | 9.2 | 215.6 | 9.75 | 2.41 | 0.63 |
| SW062 | RES02 | RES | Summer | 21 | 9.2 | 215.7 | 9.57 | 2.54 | 0.65 |
| SW065 | RES04 | RES | Summer | 20.9 | 9.3 | 214 | 10.55 | 2.3 | 0.6 |
| SW066 | RES04 | RES | Summer | 21.4 | 9.3 | 215.4 | 10.09 | 2.5 | 0.43 |
| SW067 | RES05 | RES | Summer | 21.6 | 9.3 | 216.9 | 8.95 | 2.04 | 0.61 |
| SW068 | RES05 | RES | Summer | 21.1 | 9.1 | 214.6 | 10.88 | 2.56 | 0.61 |
| SW069 | COM01 | COM | Summer | 21.8 | 9.2 | 219.5 | 11.41 | 0.26 | 0.54 |
| SW070 | COM01 | COM | Summer | 22.1 | 9.3 | 213.1 | 10.9 | 1.16 | 0.54 |
| SW071 | COM02 | COM | Summer | 21.9 | 9.1 | 217 | 10.23 | 0.03 | 0.59 |
| SW072 | COM02 | COM | Summer | 21.6 | 9.3 | 210.7 | 10.04 | 2.08 | 0.57 |
| SW073 | COM03 | COM | Summer | 22 | 8.5 | 233.9 | 9.7 | 0.92 | 0.45 |
| SW074 | COM03 | COM | Summer | 22.1 | 9.2 | 214.9 | 11.57 | 2.24 | 0.58 |
| SW082 | COM04 | COM | Summer | 21.8 | 9.2 | 214.6 | 10.11 | 1.46 | 0.53 |
| SW083 | COM04 | COM | Summer | 21.7 | 9.4 | 215.4 | 10.06 | 2.16 | 0.56 |
| SW084 | COM05 | COM | Summer | 21 | 8.9 | 232 | 9.32 | 0.02 | 0.26 |
| SW085 | COM05 | COM | Summer | 20.7 | 9.3 | 215.7 | 9.61 | 2.4 | 0.58 |
| SW087 | COM06 | COM | Summer | 21.2 | 9.4 | 217.6 | 9.87 | 2.46 | 0.59 |
| SW088 | RES01 | RES | Fall | 18.6 | 9.3 | 177 | 9.62 | 2.32 | 0.62 |
| SW089 | RES01 | RES | Fall | 18.7 | 9.6 | 176.6 | 10.11 | 2.54 | 0.61 |
| SW090 | RES02 | RES | Fall | 18.4 | 9.2 | 174.2 | 9.81 | 2.68 | 0.68 |
| SW094 | RES04 | RES | Fall | 18.8 | 9.3 | 176.5 | 9.71 | 2.38 | 0.68 |
| SW095 | RES04 | RES | Fall | 18.6 | 9.1 | 178.5 | 9.93 | 1.68 | 0.68 |
| SW096 | RES05 | RES | Fall | 18.6 | 9.1 | 177.7 | 10.71 | 2.26 | 0.74 |
| SW097 | RES05 | RES | Fall | 18.4 | 9.6 | 176.9 | 10.83 | 2.76 | 0.75 |
| SW098 | COM01 | COM | Fall | 20.7 | 9.3 | 192.8 | 9.68 | 0.08 | 0.67 |
| SW099 | COM01 | COM | Fall | 19.7 | 9.2 | 169.5 | 9.15 | 2.46 | 0.7 |
| SW100 | COM02 | COM | Fall | 20 | 9.6 | 174.1 | 10.95 | 1.58 | 0.66 |
| SW101 | COM02 | COM | Fall | 19 | 9.5 | 169.7 | 11.18 | 1.86 | 0.65 |
| SW102 | COM03 | COM | Fall | 18.2 | 9.4 | 172.7 | 11.9 | 1.2 | 0.5 |
| SW103 | COM03 | COM | Fall | 19.3 | 9.4 | 166.7 | 11.9 | 2.3 | 0.48 |
| SW112 | COM04 | COM | Fall | 18.9 | 9.3 | 166.1 | 10.1 | 2.44 | 0.65 |
| SW113 | COM05 | COM | Fall | 19.3 | 9.0 | 172.6 | 9.7 | 0.04 | 0.21 |
| SW114 | COM05 | COM | Fall | 19.2 | 9.2 | 174.1 | 9.0 | 2.22 | 0.36 |
| SW116 | COM06 | COM | Fall | 19.2 | 9.2 | 168.6 | 9.6 | 2.62 | 0.61 |
| SW117 | RES01 | RES | Fall | 19.1 | 9.4 | 224.9 | 9.9 | 2.34 | 0.52 |
| SW118 | RES01 | RES | Fall | 18.9 | 9.2 | 216.5 | 10.1 | 2.64 | 0.56 |
| SW119 | RES02 | RES | Fall | 18.4 | 9.2 | 212.9 | 10.0 | 2.84 | 0.85 |
| SW120 | RES02 | RES | Fall | 18.9 | 9.0 | 213 | 9.9 | 2.8 | 0.89 |
| SW123 | RES04 | RES | Fall | 19 | 9.2 | 215.4 | 10.0 | 2.76 | 0.76 |
| SW124 | RES04 | RES | Fall | 18.9 | 9.2 | 213.9 | 10.0 | 2.84 | 0.9 |
| SW125 | RES05 | RES | Fall | 19.1 | 9.3 | 216.2 | 10.6 | 2.54 | 0.78 |
| SW126 | RES05 | RES | Fall | 19.1 | 9.3 | 215.4 | 11.1 | 2.9 | 0.82 |
| SW127 | COM01 | COM | Fall | 20.1 | 9.2 | 216.4 | 10.8 | 0.06 | 0.49 |
| SW128 | COM01 | COM | Fall | 20.6 | 9.0 | 208.6 | 10.1 | 2.4 | 0.59 |
| SW130 | COM02 | COM | Fall | 20.9 | 9.4 | 215.1 | 11.2 | 1.98 | 0.77 |
| SW131 | COM03 | COM | Fall | 19.4 | 9.5 | 197 | 11.1 | 1.6 | 0.51 |
| SW132 | COM03 | COM | Fall | 19.3 | 9.7 | 197.1 | 11.4 | 2.9 | 0.58 |
| SW141 | COM04 | COM | Fall | 19.5 | 9.5 | 195.7 | 10.2 | 2.64 | 0.61 |
| SW143 | COM05 | COM | Fall | 18.8 | 9.4 | 197.1 | 9.7 | 2.68 | 0.61 |
| SW145 | COM06 | COM | Fall | 19.4 | 9.6 | 196 | 9.7 | 2.56 | 0.61 |
| SW152 | RES04 | RES | Fall | 19.6 | 9.5 | 172.2 | 10.3 | 2.74 | 0.57 |
| SW154 | RES05 | RES | Fall | 19.6 | 9.6 | 172.2 | 10.4 | 2.7 | 0.32 |
| SW156 | COM01 | COM | Fall | 19.7 | 9.4 | 179.1 | 10.1 | 0 | 0.12 |
| SW157 | COM01 | COM | Fall | 19.4 | 9.6 | 174.7 | 10.5 | 2.04 | 0.53 |
| SW158 | COM02 | COM | Fall | 19.9 | 9.6 | 176.1 | 10.4 | 1.48 | 0.24 |
| SW159 | COM02 | COM | Fall | 19.6 | 9.6 | 175.7 | 10.6 | 1.7 | 0.24 |
| SW160 | COM03 | COM | Fall | 22.7 | 9.5 | 178.3 | 10.8 | 0.96 | 0.14 |
| SW161 | COM03 | COM | Fall | 22.5 | 9.6 | 175.9 | 11.1 | 1.74 | 0.7 |
| SW170 | COM04 | COM | Fall | 18.3 | 9.9 | 172.1 | 9.9 | 2.78 | 0.81 |
| SW171 | COM05 | COM | Fall | 21.5 | 9.8 | 178.2 | 9.9 | 0.36 | 0.29 |
| SW172 | COM05 | COM | Fall | 21.2 | 9.5 | 179.2 | 10.4 | 2.22 | 0.68 |

**Table S2: continued**

| Sample | Nitrate  (mgN/L) | Nitrite  (mgN/L) | Total organic carbon (mgTOC/L) | Dissolved organic carbon (mgTOC/L) | Total nitrogen  (mgN/L) | Total dissolved nitrogen  (mgN/L) | Water demand  (m^3^/month) | Month |
| --- | --- | --- | --- | --- | --- | --- | --- | --- |
| SW001 | 0.6 | 0 | 0.0 | 0.0 | 51.0 | 0.0 | 933 | June |
| SW002 | 0.4 | 0 | 209.2 | 0.0 | 49.9 | 0.0 | 933 | June |
| SW003 | 0.4 | 0 | 227.5 | 0.0 | 47.7 | 0.0 | 1835 | June |
| SW004 | 0.5 | 0 | 223.9 | 0.0 | 48.7 | 0.0 | 1835 | June |
| SW007 | 0.4 | 0.003 | 229.3 | 0.0 | 51.2 | 0.0 | 1510 | June |
| SW008 | 0.4 | 0.003 | 205.0 | 0.0 | 49.7 | 0.0 | 1510 | June |
| SW009 | 0.5 | 0.004 | 221.3 | 0.0 | 50.8 | 0.0 | 1510 | June |
| SW010 | 0.4 | 0.002 | 241.3 | 0.0 | 50.0 | 0.0 | 1510 | June |
| SW011 | 0.4 | 0.007 | 158.1 | 0.0 | 57.6 | 0.0 | 71750 | June |
| SW012 | 0.4 | 0.007 | 172.2 | 0.0 | 58.9 | 0.0 | 71750 | June |
| SW014 | 0.5 | 0.006 | 204.3 | 0.0 | 52.3 | 0.0 | 71750 | June |
| SW015 | 0.4 | 0.024 | 176.3 | 173.2 | 51.8 | 54.8 | 15470 | June |
| SW016 | 0.6 | 0.013 | 182.7 | 180.0 | 52.4 | 51.3 | 15470 | June |
| SW024 | 0.4 | 0.071 | 190.0 | 0.0 | 51.3 | 0.0 | 15470 | June |
| SW025 | 0.6 | 0.093 | 190.6 | 0.0 | 51.7 | 0.0 | 15470 | June |
| SW026 | 0.7 | 0.004 | 105.4 | 0.0 | 70.8 | 0.0 | 37840 | June |
| SW027 | 0.5 | 0.001 | 176.2 | 0.0 | 48.1 | 0.0 | 37840 | June |
| SW028 | 0.4 | 0.011 | 157.9 | 0.0 | 53.0 | 0.0 | 37840 | June |
| SW029 | 0.5 | 0 | 217.0 | 0.0 | 47.0 | 0.0 | 37840 | June |
| SW030 | 0.4 | 0.004 | 750.3 | 0.0 | 49.6 | 0.0 | 1081 | July |
| SW031 | 0.3 | 0.005 | 1233.0 | 0.0 | 47.2 | 0.0 | 1081 | July |
| SW032 | 0.4 | 0 | 1055.6 | 0.0 | 46.9 | 0.0 | 1693 | July |
| SW033 | 0.4 | 0.001 | 530.0 | 0.0 | 45.7 | 0.0 | 1693 | July |
| SW037 | 0.9 | 0.002 | 1386.7 | 0.0 | 46.5 | 0.0 | 1580 | July |
| SW038 | 0.7 | 0 | 281.7 | 0.0 | 45.4 | 0.0 | 1580 | July |
| SW039 | 0.8 | 0 | 199.6 | 0.0 | 46.2 | 0.0 | 1580 | July |
| SW040 | 1.1 | 0.011 | 319.8 | 0.0 | 55.8 | 0.0 | 70950 | July |
| SW041 | 0.8 | 0.007 | 442.6 | 0.0 | 55.4 | 0.0 | 70950 | July |
| SW042 | 1 | 0.006 | 329.8 | 0.0 | 54.2 | 0.0 | 70950 | July |
| SW043 | 1.1 | 0.002 | 523.3 | 0.0 | 48.5 | 0.0 | 70950 | July |
| SW044 | 0.9 | 0.034 | 163.4 | 184.0 | 52.4 | 50.6 | 21190 | July |
| SW045 | 0.8 | 0.024 | 199.8 | 182.4 | 49.4 | 50.2 | 21190 | July |
| SW054 | 0.5 | 0.048 | 374.1 | 0.0 | 50.0 | 0.0 | 21190 | July |
| SW055 | 0.8 | 0 | 187.8 | 0.0 | 53.9 | 0.0 | 91540 | July |
| SW056 | 0.7 | 0 | 570.0 | 0.0 | 46.4 | 0.0 | 91540 | July |
| SW057 | 0.7 | 0.008 | 309.6 | 0.0 | 50.2 | 0.0 | 91540 | July |
| SW058 | 0.7 | 0 | 194.0 | 0.0 | 47.4 | 0.0 | 91540 | July |
| SW059 | 0.5 | 0.009 | 237.5 | 0.0 | 50.8 | 0.0 | 885 | August |
| SW060 | 0.5 | 0.004 | 227.7 | 0.0 | 49.4 | 0.0 | 885 | August |
| SW061 | 0.6 | 0 | 227.7 | 0.0 | 48.3 | 0.0 | 1853 | August |
| SW062 | 0.7 | 0 | 213.0 | 0.0 | 48.3 | 0.0 | 1853 | August |
| SW065 | 0.6 | 0.001 | 219.6 | 0.0 | 49.0 | 0.0 | 1680 | August |
| SW066 | 0.5 | 0 | 226.0 | 0.0 | 47.8 | 0.0 | 1680 | August |
| SW067 | 0.7 | 0.001 | 211.7 | 0.0 | 46.0 | 0.0 | 1680 | August |
| SW068 | 0.5 | 0 | 211.9 | 0.0 | 47.1 | 0.0 | 1680 | August |
| SW069 | 0.7 | 0.021 | 286.8 | 0.0 | 56.5 | 0.0 | 76770 | August |
| SW070 | 0.5 | 0.017 | 347.8 | 0.0 | 53.6 | 0.0 | 76770 | August |
| SW071 | 0.6 | 0.023 | 199.8 | 0.0 | 56.6 | 0.0 | 76770 | August |
| SW072 | 0.6 | 0.015 | 225.6 | 0.0 | 49.4 | 0.0 | 76770 | August |
| SW073 | 0.6 | 0.054 | 185.5 | 185.5 | 52.0 | 52.8 | 39620 | August |
| SW074 | 0.7 | 0.023 | 170.6 | 201.3 | 48.7 | 48.8 | 39620 | August |
| SW082 | 0.4 | 0.022 | 479.5 | 0.0 | 84.9 | 0.0 | 39620 | August |
| SW083 | 0.5 | 0.021 | 356.1 | 0.0 | 56.9 | 0.0 | 39620 | August |
| SW084 | 0.9 | 0.011 | 206.3 | 0.0 | 58.4 | 0.0 | 83840 | August |
| SW085 | 0.4 | 0.009 | 286.6 | 0.0 | 71.2 | 0.0 | 83840 | August |
| SW087 | 0.5 | 0.006 | 339.9 | 0.0 | 85.3 | 0.0 | 83840 | August |
| SW088 | 0.7 | 0.031 | 564.0 | 0.0 | 55.6 | 0.0 | 748 | September |
| SW089 | 0.7 | 0.033 | 538.4 | 0.0 | 76.2 | 0.0 | 748 | September |
| SW090 | 0.7 | 0 | 648.0 | 0.0 | 0.0 | 0.0 | 2253 | September |
| SW094 | 0.8 | 0.004 | 563.1 | 0.0 | 78.9 | 0.0 | 1530 | September |
| SW095 | 0.8 | 0.004 | 0.0 | 0.0 | 58.9 | 0.0 | 1530 | September |
| SW096 | 0.8 | 0.016 | 415.6 | 0.0 | 0.0 | 0.0 | 1530 | September |
| SW097 | 0.7 | 0.004 | 0.0 | 0.0 | 80.4 | 0.0 | 1530 | September |
| SW098 | 0.03 | 0.03 | 440.3 | 0.0 | 74.4 | 0.0 | 65850 | September |
| SW099 | 0.5 | 0.027 | 0.0 | 0.0 | 95.3 | 0.0 | 65850 | September |
| SW100 | 0.7 | 0.042 | 494.5 | 0.0 | 0.0 | 0.0 | 65850 | September |
| SW101 | 0.7 | 0.029 | 337.5 | 0.0 | 95.5 | 0.0 | 65850 | September |
| SW102 | 1 | 0.23 | 171.4 | 168.1 | 54.7 | 53.9 | 45240 | September |
| SW103 | 0.9 | 0.13 | 176.3 | 171.5 | 51.5 | 51.9 | 45240 | September |
| SW112 | 0.7 | 0.17 | 319.4 | 0.0 | 51.1 | 0.0 | 45240 | September |
| SW113 | 1.2 | -0.01 | 128.9 | 0.0 | 55.5 | 0.0 | 35910 | September |
| SW114 | 0.7 | 0.24 | 193.3 | 0.0 | 50.6 | 0.0 | 35910 | September |
| SW116 | 0.9 | 0.21 | 192.7 | 0.0 | 50.2 | 0.0 | 35910 | September |
| SW117 | 0.7 | 0.043 | 240.0 | 0.0 | 53.6 | 0.0 | 757 | October |
| SW118 | 0.7 | 0.032 | 202.1 | 0.0 | 49.5 | 0.0 | 757 | October |
| SW119 | 0.8 | 0.003 | 237.2 | 0.0 | 47.1 | 0.0 | 2230 | October |
| SW120 | 0.7 | 0.004 | 218.4 | 0.0 | 49.3 | 0.0 | 2230 | October |
| SW123 | 0.6 | 0.006 | 200.6 | 0.0 | 47.8 | 0.0 | 1580 | October |
| SW124 | 0.006 | 0.85 | 212.5 | 0.0 | 46.9 | 0.0 | 1580 | October |
| SW125 | 0.7 | 0.006 | 219.1 | 0.0 | 48.7 | 0.0 | 1580 | October |
| SW126 | 0.8 | 0.009 | 0.0 | 0.0 | 47.9 | 0.0 | 1580 | October |
| SW127 | 0.8 | 0.006 | 162.6 | 0.0 | 58.2 | 0.0 | 46000 | October |
| SW128 | 0.8 | 0.035 | 196.5 | 0.0 | 50.9 | 0.0 | 46000 | October |
| SW130 | 0.6 | 0.025 | 255.0 | 0.0 | 54.0 | 0.0 | 46000 | October |
| SW131 | 0.6 | 0.049 | 0.0 | 176.5 | 56.2 | 52.6 | 48020 | October |
| SW132 | 0.6 | 0.032 | 167.0 | 189.7 | 50.0 | 52.8 | 48020 | October |
| SW141 | 0.5 | 0.02 | 197.1 | 0.0 | 49.2 | 0.0 | 48020 | October |
| SW143 | 0.7 | 0.023 | 198.6 | 0.0 | 50.8 | 0.0 | 37210 | October |
| SW145 | 0.8 | 0.022 | 277.5 | 0.0 | 48.1 | 0.0 | 37210 | October |
| SW152 | 0.6 | 0.002 | 213.0 | 0.0 | 44.6 | 0.0 | 1890 | November |
| SW154 | 0.7 | 0.002 | 222.1 | 0.0 | 45.0 | 0.0 | 1890 | November |
| SW156 | 0.6 | 0.007 | 158.1 | 0.0 | 55.1 | 0.0 | 25700 | November |
| SW157 | 1 | 0.015 | 163.5 | 0.0 | 45.9 | 0.0 | 25700 | November |
| SW158 | 0.7 | 0.011 | 171.7 | 0.0 | 48.7 | 0.0 | 25700 | November |
| SW159 | 0.6 | 0.008 | 156.9 | 0.0 | 47.6 | 0.0 | 25700 | November |
| SW160 | 1 | 0.011 | 140.0 | 0.0 | 55.4 | 53.1 | 42710 | November |
| SW161 | 0.9 | 0.01 | 144.1 | 154.9 | 46.0 | 48.1 | 42710 | November |
| SW170 | 0.7 | 0.004 | 166.7 | 0.0 | 43.0 | 0.0 | 42710 | November |
| SW171 | 0.9 | 0.048 | 153.0 | 0.0 | 51.2 | 0.0 | 41180 | November |
| SW172 | 0.8 | 0.01 | 0.0 | 0.0 | 50.1 | 0.0 | 41180 | November |

**Table S3: Read abundance in commercial and residential samples showing for each OTU with associated taxonomy classification. P-value were adjusted with the Benjamin and Hochberg method. P-value < 0.05 were considered significant differential abundance.**

| **Taxa** | **Read abundance in RES (commercial) samples** | **Read abundance in COM (residential) samples** | **P-value adjusted** |
| --- | --- | --- | --- |
| OTU1979 Curvibacter | 16 | 58679 | 5.9869E-74 |
| OTU4095 Qipengyuania | 19 | 16010 | 6.5563E-49 |
| OTU4541 Porphyrobacter | 100 | 23646 | 1.8857E-40 |
| OTU2099 Sediminibacterium | 2 | 11484 | 6.1759E-39 |
| OTU2466 Gemmatimonas | 50 | 9845 | 1.5938E-36 |
| OTU1874 Acidovorax delafieldii | 155 | 13230 | 1.2633E-26 |
| OTU4815 Phreatobacter oligotrophus | 137 | 39025 | 8.5691E-26 |
| OTU1749 TRA3-20 | 4 | 15405 | 8.0902E-23 |
| OTU1282 UBA12409 | 2 | 5894 | 7.0116E-19 |
| OTU2008 Sediminibacterium | 10 | 8637 | 7.1376E-16 |
| OTU1992 Sediminibacterium | 25 | 12157 | 5.0524E-15 |
| OTU5231 Brevundimonas bacteroides | 53 | 6697 | 4.8516E-12 |
| OTU4317 Blastomonas | 36 | 5205 | 5.5907E-11 |
| OTU4699 Novosphingobium | 868 | 8678 | 3.2074E-08 |
| OTU1845 Herminiimonas | 248 | 6700 | 3.7702E-08 |
| OTU4423 Sphingomonas | 1167 | 15644 | 3.8039E-08 |
| OTU2064 Hydrogenophaga | 54 | 20213 | 4.9939E-08 |
| OTU1277 Nitrospira | 272 | 5389 | 8.56E-08 |
| OTU2912 Mycobacterium frederiksbergense | 7654 | 1977 | 8.2251E-06 |
| OTU1744 Simplicispira | 4259 | 1754 | 0.00020352 |
| OTU2472 Paucibacter | 3884 | 1684 | 0.00025273 |
| OTU1689 TRA3-20 | 34 | 7783 | 0.00026656 |
| OTU3398 Flavobacterium | 4144 | 1697 | 0.00043248 |
| OTU1492 Rhodoferax | 6879 | 3379 | 0.0010596 |
| OTU1683 Duganella | 3718 | 1583 | 0.00152624 |
| OTU1486 Neisseriaceae | 59068 | 29218 | 0.00183469 |
| OTU4270 Sphingomonas | 4820 | 14937 | 0.00196045 |
| OTU4225 Sphingomonas | 25854 | 15909 | 0.00289771 |
| OTU1667 Undibacterium | 3936 | 2166 | 0.00355915 |
| OTU1867 Duganella | 3741 | 1572 | 0.00400826 |
| OTU1487 Pseudomonas | 12745 | 7633 | 0.00689642 |
| OTU2092 Rugamonas | 5214 | 2963 | 0.00698765 |
| OTU4694 Methylobacterium-Methylorubrum | 5944 | 17264 | 0.01125242 |
| OTU1840 Undibacterium | 9122 | 5850 | 0.01992029 |
| OTU2502 Hydrogenophaga | 6447 | 3496 | 0.02818611 |
| OTU1838 Duganella | 7215 | 5052 | 0.04196739 |
| OTU1765 Deefgea | 3495 | 2255 | 0.04347225 |
| OTU1494 Pseudomonas peli | 7580 | 5600 | 0.05559141 |
| OTU4366 Magnetospirillum | 3563 | 1575 | 0.05559141 |
| OTU3945 MD3-55 | 5955 | 4131 | 0.05863204 |
| OTU4564 Methylobacterium-Methylorubrum rhodesianum | 7635 | 1244 | 0.07475509 |
| OTU3203 Campylobacterales | 4036 | 2473 | 0.09435376 |
| OTU1491 Pseudomonas | 3998 | 3168 | 0.09735811 |
| OTU4431 Telmatospirillum | 6834 | 3654 | 0.11560097 |
| OTU1376 Nitrosomonas oligotropha | 1512 | 3629 | 0.14672868 |
| OTU1010 Nitrospira | 1939 | 3820 | 0.20780288 |
| OTU1637 Paludibacterium paludis | 5406 | 4708 | 0.21662227 |
| OTU4580 Methylobacterium-Methylorubrum adhaesivum | 9082 | 8963 | 0.50177286 |
| OTU1377 Methylotenera | 8545 | 11287 | 0.87167283 |

**Table S4: Bin id, bin relative abundance, top taxa, bin contribution proportions to each ecological process for samples collected from COM. Ecological processes include homogeneous selection (HoS), heterogeneous selection (HeS), dispersal limitation (DL), homogenizing dispersal (HD), and drift (DR).**

Bin 1: Proteobacteria; Gammaproteobacteria; Burkholderiales; Oxalobacteraceae; Duganella

Bin 2: Proteobacteria; Gammaproteobacteria; Burkholderiales; Oxalobacteraceae; Rugamonas

Bin 3: Proteobacteria; Gammaproteobacteria; Burkholderiales; Oxalobacteraceae; Duganella

Bin 4: Proteobacteria; Gammaproteobacteria; Burkholderiales; Oxalobacteraceae; Herminiimonas

Bin 5: Proteobacteria; Alphaproteobacteria; Rhizobiales; Rhizobiales Incertae Sedis; Phreatobacter oligotrophus

Bin 6: Proteobacteria; Gammaproteobacteria; Burkholderiales; Neisseriaceae

Bin 7: Proteobacteria; Gammaproteobacteria; Burkholderiales; Comamonadaceae; Curvibacter delafieldii

Bin 8: Proteobacteria; Gammaproteobacteria; Burkholderiales; Comamonadaceae; Paucibacter

| *Bin_id* | | Bin1 | Bin2 | Bin3 | Bin4 | Bin5 | Bin6 | Bin7 | Bin8 |
| --- | --- | --- | --- | --- | --- | --- | --- | --- | --- |
| *Bin_relative_abundance* | | 00.0078 | 00.0076 | 00.0125 | 00.0344 | 00.6697 | 00.0690 | 00.1860 | 00.0130 |
| *Dominant_Process* | *June* | DL | DR | DL | DR | DR | DL | DR | DL |
|  | *July* | DL | DR | DL | DR | DR | DL | DR | DR |
|  | *August* | DL | DL | DL | HeS | DL | DR | DL | DL |
|  | *September* | DL | DL | DR | DR | DL | DR | DR | DL |
|  | *October* | DL | DL | DR | DR | DR | DR | DR | DR |
|  | *November* | DR | DR | DR | DR | DR | HD | DR | DR |
| *Bin_contribution_to_process_June* | *HeS* | 0.000000 | 0.000000 | 0.000000 | 0.000000 | 0.000000 | 0.000000 | 0.003859 | 0.000000 |
|  | *HoS* | 0.001664 | 0.000000 | 0.000000 | 0.000000 | 0.118068 | 0.000000 | 0.000000 | 0.000000 |
|  | *DL* | 0.007151 | 0.000672 | 0.008453 | 0.000000 | 0.266917 | 0.063132 | 0.023088 | 0.005916 |
|  | *HD* | 0.000000 | 0.000007 | 0.000000 | 0.001010 | 0.021910 | 0.001554 | 0.006198 | 0.000051 |
|  | *DR* | 0.000386 | 0.001093 | 0.004327 | 0.011112 | 0.288102 | 0.039897 | 0.121373 | 0.004060 |
| *Bin_contribution_to_process_July* | *HeS* | 0.000000 | 0.000056 | 0.000000 | 0.000814 | 0.035899 | 0.000000 | 0.018656 | 0.000851 |
|  | *HoS* | 0.001344 | 0.000000 | 0.001008 | 0.000675 | 0.000000 | 0.000000 | 0.000000 | 0.000313 |
|  | *DL* | 0.007907 | 0.000479 | 0.005024 | 0.003282 | 0.208954 | 0.083795 | 0.067121 | 0.004674 |
|  | *HD* | 0.000000 | 0.000005 | 0.000000 | 0.000871 | 0.013875 | 0.000569 | 0.000000 | 0.000044 |
|  | *DR* | 0.000419 | 0.002209 | 0.004843 | 0.012710 | 0.356747 | 0.049272 | 0.112351 | 0.005234 |
| *Bin_contribution_to_process_August* | *HeS* | 0.000173 | 0.001705 | 0.002011 | 0.019062 | 0.067361 | 0.000000 | 0.006743 | 0.000000 |
|  | *HoS* | 0.000000 | 0.000314 | 0.005362 | 0.000000 | 0.015711 | 0.000000 | 0.000000 | 0.000000 |
|  | *DL* | 0.008773 | 0.016341 | 0.008105 | 0.013588 | 0.363993 | 0.000000 | 0.119858 | 0.014193 |
|  | *HD* | 0.000000 | 0.000000 | 0.000000 | 0.000345 | 0.039676 | 0.010492 | 0.003304 | 0.000010 |
|  | *DR* | 0.000607 | 0.002419 | 0.003340 | 0.015439 | 0.182981 | 0.014221 | 0.060154 | 0.003718 |
| *Bin_contribution_to_proces_September* | *HeS* | 0.000000 | 0.000000 | 0.000000 | 0.009763 | 0.000000 | 0.000000 | 0.000000 | 0.000484 |
|  | *HoS* | 0.000000 | 0.000000 | 0.003359 | 0.000000 | 0.031372 | 0.000000 | 0.000000 | 0.000000 |
|  | *DL* | 0.006874 | 0.004646 | 0.004175 | 0.013083 | 0.365577 | 0.000000 | 0.013518 | 0.011001 |
|  | *HD* | 0.000000 | 0.000000 | 0.000000 | 0.001091 | 0.027824 | 0.005069 | 0.001478 | 0.000012 |
|  | *DR* | 0.001186 | 0.002712 | 0.007390 | 0.017350 | 0.265649 | 0.055069 | 0.145253 | 0.006063 |
| *Bin_contribution_to_process_October* | *HeS* | 0.000000 | 0.000000 | 0.000000 | 0.004997 | 0.000000 | 0.000000 | 0.000000 | 0.000925 |
|  | *HoS* | 0.000000 | 0.000000 | 0.001372 | 0.002923 | 0.000000 | 0.000000 | 0.000000 | 0.000000 |
|  | *DL* | 0.006418 | 0.005625 | 0.004908 | 0.020095 | 0.145206 | 0.000000 | 0.066058 | 0.007615 |
|  | *HD* | 0.000000 | 0.000000 | 0.000000 | 0.000119 | 0.047006 | 0.010131 | 0.001794 | 0.000000 |
|  | *DR* | 0.002448 | 0.002568 | 0.008286 | 0.020257 | 0.446063 | 0.066636 | 0.118129 | 0.010419 |
| *Bin_contribution_to_process_November* | *HeS* | 0.000000 | 0.000000 | 0.000000 | 0.008150 | 0.019561 | 0.000000 | 0.000000 | 0.000000 |
|  | *HoS* | 0.000000 | 0.000709 | 0.000538 | 0.000000 | 0.000000 | 0.000334 | 0.000000 | 0.000000 |
|  | *DL* | 0.000000 | 0.000000 | 0.000000 | 0.010032 | 0.176538 | 0.000000 | 0.063816 | 0.000000 |
|  | *HD* | 0.000000 | 0.000000 | 0.000052 | 0.000324 | 0.000000 | 0.003330 | 0.000000 | 0.000125 |
|  | *DR* | 0.000558 | 0.003514 | 0.001095 | 0.025575 | 0.513697 | 0.000000 | 0.169803 | 0.002247 |

**Table S5: Bin id, bin relative abundance, bin contribution proportions to each ecological process for samples collected from RES. Ecological processes include homogeneous selection (HoS), heterogeneous selection (HeS), dispersal limitation (DL), homogenizing dispersal (HD), and drift (DR).**

Bin 1: Proteobacteria; Gammaproteobacteria; Burkholderiales; Neisseriaceae; Duganella paludism

Bin 2: Proteobacteria; Gammaproteobacteria; Pseudomonadales; Pseudomonadaceae; Pseudomonas anguilliseptica

Bin 3: Proteobacteria; Alphaproteobacteria; Rhizobiales; Beijerinckiaceae; Methylobacterium-Methylorubrum adhaesivum

Bin 4: Proteobacteria; Alphaproteobacteria; Sphingomonadales; Sphingomonadaceae; Sphingomonas natatoria

Bin 5: Actinobacteriota; Actinobacteria; Corynebacteriales; Mycobacteriaceae; Mycobacterium frederiksbergense

| *Bin_id* | | Bin1 | Bin2 | Bin3 | Bin4 | Bin5 |
| --- | --- | --- | --- | --- | --- | --- |
| *Bin_relative_abundance* | | 0.574 | 0.100 | 0.137 | 0.133 | 0.054 |
| *Dominant_Process* | *June* | DR | DR | DL | DR | HeS |
|  | *July* | DR | DR | DR | DR | DL |
|  | *August* | HeS | DR | DR | DR | DR |
|  | *September* | HeS | DR | DR | DR | DL |
|  | *October* | HeS | DR | DR | DR | DL |
|  | *November* | HeS | DR | DR | DR | DL |
| *Bin_contribution_to_process_June* | *HeS* | 0.000000 | 0.002645 | 0.000000 | 0.000000 | 0.052658 |
|  | *HoS* | 0.000000 | 0.000000 | 0.000585 | 0.020258 | 0.001571 |
|  | *DL* | 0.000000 | 0.000000 | 0.048819 | 0.000000 | 0.009985 |
|  | *HD* | 0.137439 | 0.010684 | 0.002072 | 0.001380 | 0.003417 |
|  | *DR* | 0.584808 | 0.047511 | 0.034373 | 0.041795 | 0.000000 |
| *Bin_contribution_to_process_July* | *HeS* | 0.000000 | 0.000000 | 0.000000 | 0.000000 | 0.025879 |
|  | *HoS* | 0.000000 | 0.000000 | 0.000000 | 0.000000 | 0.000000 |
|  | *DL* | 0.000000 | 0.000000 | 0.014433 | 0.000000 | 0.026768 |
|  | *HD* | 0.185003 | 0.004797 | 0.000000 | 0.000632 | 0.000000 |
|  | *DR* | 0.492979 | 0.075607 | 0.085722 | 0.088180 | 0.000000 |
| *Bin_contribution_to_process_August* | *HeS* | 0.483148 | 0.039341 | 0.000000 | 0.000000 | 0.000963 |
|  | *HoS* | 0.000000 | 0.000000 | 0.000000 | 0.000000 | 0.000000 |
|  | *DL* | 0.000000 | 0.038771 | 0.057403 | 0.016416 | 0.000000 |
|  | *HD* | 0.000000 | 0.000000 | 0.000000 | 0.000000 | 0.002909 |
|  | *DR* | 0.000000 | 0.078419 | 0.113682 | 0.148313 | 0.020635 |
| *Bin_contribution_to_proces_September* | *HeS* | 0.475102 | 0.000000 | 0.000000 | 0.000000 | 0.000000 |
|  | *HoS* | 0.000000 | 0.000000 | 0.000000 | 0.000000 | 0.000000 |
|  | *DL* | 0.000000 | 0.000000 | 0.088252 | 0.061922 | 0.028053 |
|  | *HD* | 0.000000 | 0.012123 | 0.008016 | 0.000000 | 0.002668 |
|  | *DR* | 0.000000 | 0.078802 | 0.115945 | 0.109634 | 0.019484 |
| *Bin_contribution_to_process_October* | *HeS* | 0.369409 | 0.000000 | 0.000000 | 0.000000 | 0.000000 |
|  | *HoS* | 0.000000 | 0.000000 | 0.000000 | 0.000000 | 0.000000 |
|  | *DL* | 0.000000 | 0.000000 | 0.057443 | 0.015092 | 0.040661 |
|  | *HD* | 0.036760 | 0.003701 | 0.000000 | 0.000000 | 0.000879 |
|  | *DR* | 0.112437 | 0.102465 | 0.082066 | 0.154488 | 0.024600 |
| *Bin_contribution_to_process_November* | *HeS* | 0.557714 | 0.000000 | 0.000000 | 0.000000 | 0.000000 |
|  | *HoS* | 0.000000 | 0.000000 | 0.000000 | 0.000000 | 0.000000 |
|  | *DL* | 0.000000 | 0.000000 | 0.000000 | 0.000000 | 0.088334 |
|  | *HD* | 0.000000 | 0.000000 | 0.000000 | 0.000000 | 0.000000 |
|  | *DR* | 0.000000 | 0.114955 | 0.072287 | 0.166710 | 0.000000 |

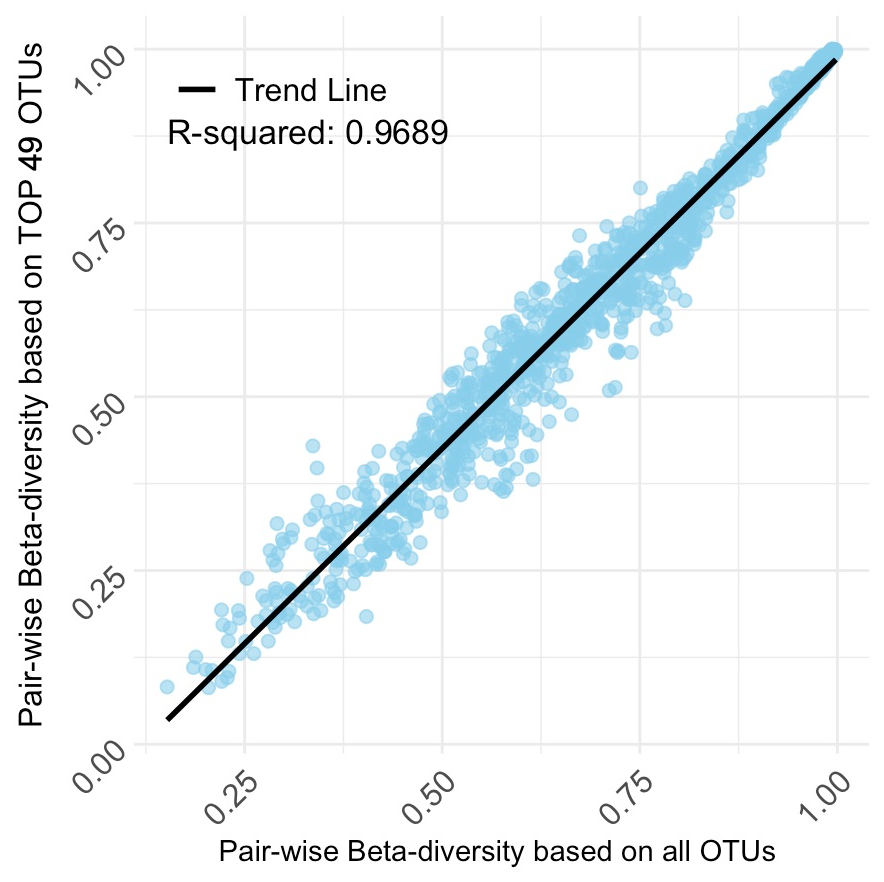

**Figure S1. Correlation between pairwise Bray-Curtis dissimilarities based on all OTUs (n = 5395), and 2) OTUs with global abundance over 5000 (n = 49).**

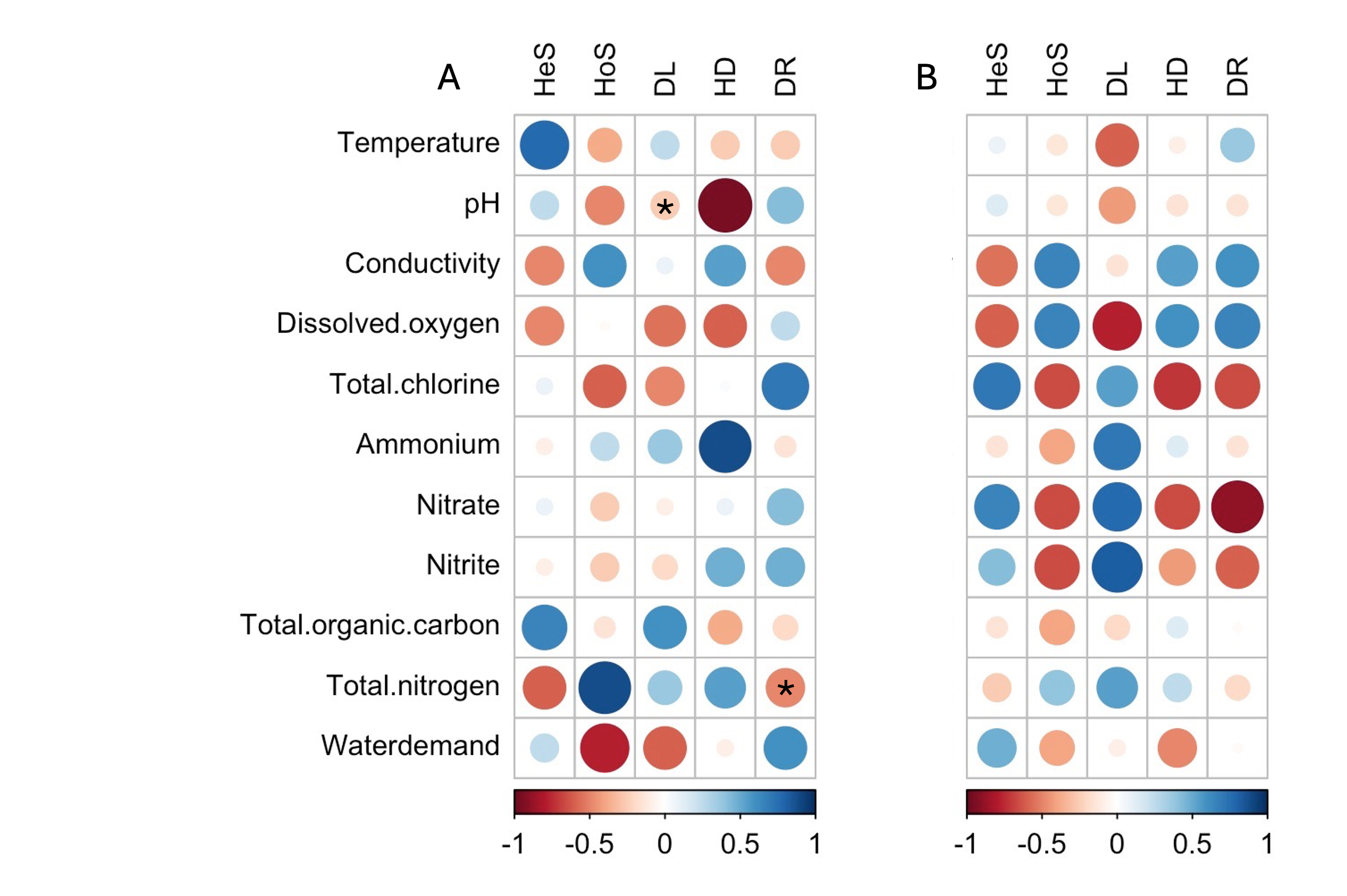

**Figure S2. Pearson correlation between ecological processes and measured environmental variables in COM (A) and RES (B). *: p (pH) = 0.016, p (total nitrogen) = 0.033. In RES, all correlations displayed non-significant p values (p > 0.05). Ecological processes include homogeneous selection (HoS), heterogeneous selection (HeS), dispersal limitation (DL), homogenizing dispersal (HD), and drift (DR).**
